# Translation of a small upstream open reading frame functions as a rheostat for the regulation of *lin-41* by the Let-7 microRNA in *Caenorhabditis elegans*

**DOI:** 10.64898/2026.09.22.750518

**Authors:** Caroline A. Spike, Micah D. Gearhart, Katherine M. Walstrom, Mark E. Zweifel, Naomi Courtemanche, David Greenstein

## Abstract

MicroRNAs have been likened to the “dark matter” of eukaryotic genomes, reflecting their pervasive regulatory influence. MicroRNAs were first identified through genetic studies of developmental timing in the nematode *Caenorhabditis elegans*. Let-7 was the first microRNA recognized to be broadly conserved. The principal target of Let-7 in the developmental timing pathway is the TRIM-NHL RNA-binding protein LIN-41. During the L4 larval stage, Let-7 represses *lin-41* translation by binding to two Let-7 complementary sites in the *lin-41* 3’UTR. Despite the importance of microRNA-based translational regulation, the underlying molecular mechanisms are incompletely understood. Through genetic analysis, we discovered an unrecognized feature of the mechanism by which Let-7 controls *lin-41* translation. This mechanism requires a 5’-regulatory exon containing a seven-amino acid upstream open reading frame (uORF) and conserved sequence elements. Genome editing indicates that the specific uORF amino acid sequence itself is not important. Our data suggest that uORF translation and 5’UTR structure limit initiation at the downstream *lin-41* start codon, enabling tight control by Let-7. Without this mechanism, the Let-7 microRNA is unable to properly regulate *lin-41* to enable proper development.

**Summary statement:** This study reports the discovery that tight control of *lin-41* translation by the Let-7 microRNA in *C. elegans* requires a 5’-regulatory exon containing a small upstream open reading frame.

## INTRODUCTION

Genetic analysis of cell growth, development and behavior in model organisms led to the identification of several developmental control genes (Meneely and Herman, 1979; Chalfie et al., 1981; Ambros and Horvitz, 1984; Weigel et al., 2000; Hipfner et al., 2002; Chang et al., 2003) that were later shown to encode microRNAs (Lee et al., 1993; Reinhart et al., 2000; Palatnik et al., 2003; Brennecke et al., 2003; Johnston and Hobert, 2003). MicroRNAs are small non-coding RNAs of approximately 22 nucleotides that are widespread in most eukaryotes (Lagos-Quintana et al., 2001; Lau et al., 2001; Lee and Ambros, 2001; Reinhart et al., 2002).

Imperfect base pairing between microRNAs and complementary sequences, predominantly within the 3’UTRs of their target genes, regulates gene expression at a post-transcriptional level (Wightman et al., 1993; Ha et al., 1996; Olsen and Ambros, 1999). In situations in which there is perfect complementarity between the microRNA and its mRNA target, mRNA degradation occurs as the predominant regulatory effect (Llave et al., 2002; Rhoades et al., 2002; Yekta et al., 2004). MicroRNAs recruit a large multi-subunit complex containing Argonaute proteins, called the microRNA-induced silencing complex (miRISC), to post-transcriptionally silence target genes (Jonas and Izaurralde, 2015; Duchaine and Fabian, 2019; Naeli et al., 2022). Molecular, biochemical, structural and single-molecule studies have converged on the viewpoint that miRISC primarily inhibits gene expression through repression of cap-dependent translation initiation with mRNA decay occurring subsequently (Bagga et al., 2005; Pillai et al., 2005; Chendrimada et al., 2007; Mathonnet et al., 2007; Eulalio et al., 2008; Bazzini et al., 2012; Djuranovic et al., 2012; Meijer et al., 2013; Cialek et al., 2022); however, the underlying molecular details and differences between experimental systems have not been fully resolved. In this study, we have taken a developmental genetics approach to address the mechanism by which the Let-7 microRNA post-transcriptionally regulates its target *lin-41* in the heterochronic gene regulatory pathway in the nematode *Caenorhabditis elegans*.

The heterochronic gene regulatory pathway controls developmental timing in *C. elegans*. Mutations in this pathway transform cell fates in time such that cells inappropriately adopt fates that are characteristic of earlier or later developmental stages (Ambros and Horvitz, 1984; Rougvie and Moss, 2013; Antebi, 2013). For example, *let-7* mutations affect the terminal differentiation of epithelial cells such that in the adult stage, these cells fail to differentiate properly and continue to divide as they do in the L4 larval stage (Reinhart et al., 2000; Slack et al., 2000). *let-7* null mutants exhibit lethality at the early adult stage with the animals bursting with their intestines extruded through the vulval opening (Meneely and Herman, 1979). The principal target of the Let-7 microRNA is the tripartite motif (TRIM)-NHL (NCL-1, HT2A and LIN-41) RNA-binding protein LIN-41 (Fig. 1A; Reinhart et al., 2000; Slack et al., 2000; Ecsedi et al., 2015; Aeschimann et al., 2019). The Let-7 microRNA represses LIN-41 translation by binding to two Let-7 complementary sites (LCSs) in the *lin-41* 3’UTR (Reinhart et al., 2000; Slack et al., 2000; Ecsedi et al., 2015). Over-expression of LIN-41 produces a lethal phenotype that phenocopies a *let-7* null mutation (Reinhart et al., 2000). In turn, LIN-41 represses the translation of the zinc-finger transcription factor LIN-29 through direct binding of sequences in the *lin-29a* 5’UTR (Fig. 1A; Aeschimann et al., 2017; Kumari et al., 2018). LIN-29 is expressed in epidermal cells in the L4 larval stage and promotes their terminal differentiation (Liu et al., 1995; Rougvie and Ambros, 1995; Bettinger et al., 1996). LIN-41 is a conserved protein (TRIM71) that regulates stem cell fate in mammals (Maller Schulman et al., 2008; Chang et al., 2012; Kwon et al., 2013; Worringer et al., 2014; Cuevas et al., 2015; Mitschka et al., 2015; Li et al., 2019); pathogenic LIN-41/Trim71 variants cause congenital hydrocephalus (Furey et al., 2018; Liu et al., 2023a,b; Duy et al., 2024) and are linked to germ cell loss and male infertility in mice and humans (Du et al., 2020; Torres-Fernández et al., 2021). Regulation of LIN-41 by *let-7* is also conserved in mammals (Pasquinelli et al., 2000; Worringer et al., 2014).

**Fig. 1.**
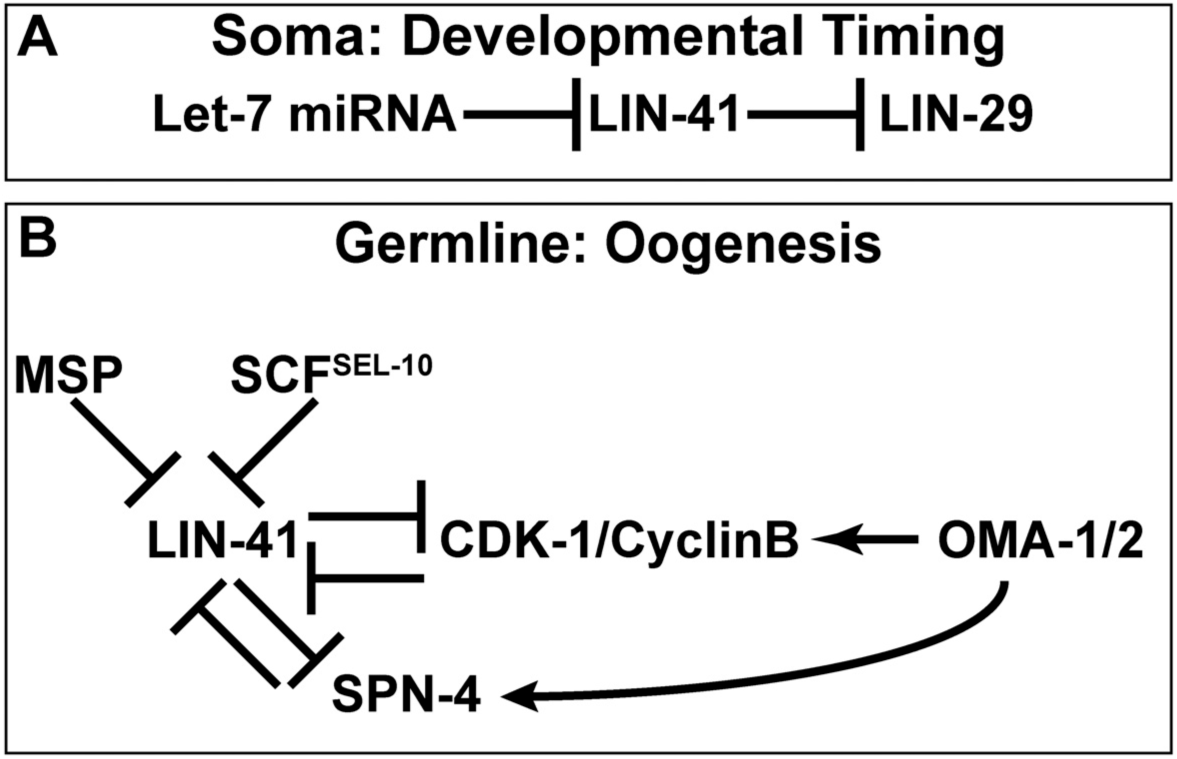
Regulation of LIN-41 in the soma and germline. (A) Regulation of LIN-41 translation in the soma. (B) Regulation of LIN-41 activity in the germline.

In addition to its role in the heterochronic gene regulatory pathway in the soma, *lin-41* is required for the extended meiotic prophase of *C. elegans* oocytes (Spike et al., 2014a,b; Tocchini et al, 2014). In *lin-41* null mutants, developing oocytes prematurely activate the CDK-1 cyclin-dependent kinase and enter M-phase, resulting in sterility (Fig. 1B). In contrast the TIS11 zinc-finger RNA-binding proteins OMA-1 and OMA-2 (OMA proteins) promote CDK-1 activation and M-phase entry of the most proximal oocytes (Detwiler et al., 2001), which occurs in response to the major sperm protein (MSP) oocyte meiotic maturation signal (Miller et al., 2001). LIN-41 is the earliest known marker of oogenesis in *C. elegans*, and it is abundantly expressed in developing oocytes where it localizes to ∼40S ribonucleoprotein particles, associates with ∼1000 mRNAs and controls translation to regulate oocyte differentiation and cell cycle progression (Spike et al., 2014a,b; Tocchini et al., 2014; Tsukamoto et al., 2017). Distinct mechanisms regulate *lin-41* activity in the germline and soma (Fig. 1). Whereas *let-7* regulates *lin-41* in the soma (Fig. 1A), the Let-7 microRNA is not expressed in the *C. elegans* germline (Lau et al., 2001) and LIN-41 protein levels were not increased in adult-stage germlines by *let-7* mutations (Spike et al., 2014a). Instead, multiple mechanisms, operating at the protein and mRNA levels control *lin-41* activity in the germline and the early embryo (Fig. 1B). Upon the onset of meiotic maturation, CDK-1 activation triggers LIN-41 protein degradation by the SCF^SEL-10^ E3 ubiquitin ligase (Spike et al., 2018). *lin-41* mRNA is subsequently degraded by a SPN-4-dependent mechanism after fertilization (Spike et al., 2026). In genetic studies of *C. elegans* oogenesis, we unexpectedly recovered novel *lin-41* mutant alleles that impact the regulation of *lin-41* by *let-7* in the soma. Analysis of these mutant alleles and subsequent genetic analyses has led us to propose a new model for the regulation of *lin-41* translation by *let-7*.

In this study, we show that Let-7 microRNA control of *lin-41* translation in somatic cells requires a 5’-regulatory exon containing a seven-amino acid upstream open reading frame (uORF). The specific *lin-41* uORF amino acid sequence itself is not important for Let-7 regulation. Rather, our results support a model in which uORF translation and 5’UTR sequences limit translation initiation at the downstream *lin-41* start codon, enabling tight control by Let-7. The engagement of uORFs with scanning ribosomes has been shown to regulate translation of the downstream main open reading frame in many contexts, including the regulation of translation in response to starvation or stress (Zhang et al., 2019; Dever et al., 2023). uORFs are found in 40-50% of human 5’UTRs (Calvo et al., 2009), potentially reflecting widespread use of microRNA regulation. uORF-dependent regulation of microRNA activity may impose switch-like control on genetic pathways organized around dosage-sensitive microRNA targets.

## RESULTS

### Novel *lin-41* reduction-of-function mutations map to an upstream regulatory exon

Prior genetic screens isolated more than 30 *lin-41* loss-of-function (lf) and reduction-of-function (rf) mutant alleles as dominant suppressors of the *let-7(n2853*ts) mutation (Slack et al., 2000; Spike et al., 2014a). Most of these *lin-41* mutations map to coding exons 1-15 or affect splice junctions (Slack et al., 2000; Spike et al., 2014a; Fig. 2A). Molecular lesions were not previously found for three viable and fertile alleles, *tn1485*, *tn1493* and *tn1503* and one sterile allele, *tn1494*. We reanalyzed these *lin-41* mutant alleles and found that *lin-41(tn1494)* resulted from a P942S mutation affecting the third NHL repeat. In addition, whole genome sequencing indicated that *lin-41(tn1485)* is associated with a complex rearrangement (*tnC1*) in which the *lin-41* locus was duplicated and some sequences from the X chromosome appeared to be inserted on chromosome I. We were unable to find the molecular lesions associated with the soma-specific *lin-41(*rf) alleles, *tn1493* and *tn1503*, in the *lin-41* coding region, splice junctions or 3’UTR. We examined transcriptomic data (Tourasse et al., 2017; Sternberg et al., 2024) and the structure of *lin-41* transcripts using reverse-transcription PCR (RT-PCR) and sequencing. These analyses demonstrated that *lin-41* is encoded by two SL1-transpliced transcripts: a long transcript that includes a 179 nucleotide 5’-exon (exon 0), and a short transcript with a 38 nucleotide 5’-UTR, including the 22 nucleotide SL1 leader sequence (Figs 2A and S1). Both transcripts use the same initiator methionine, consistent with the generation of functional proteins by fusing GFP and mScarlet sequences to the amino-terminus of LIN-41 (Spike et al., 2014a, 2022).

**Fig. 2.**
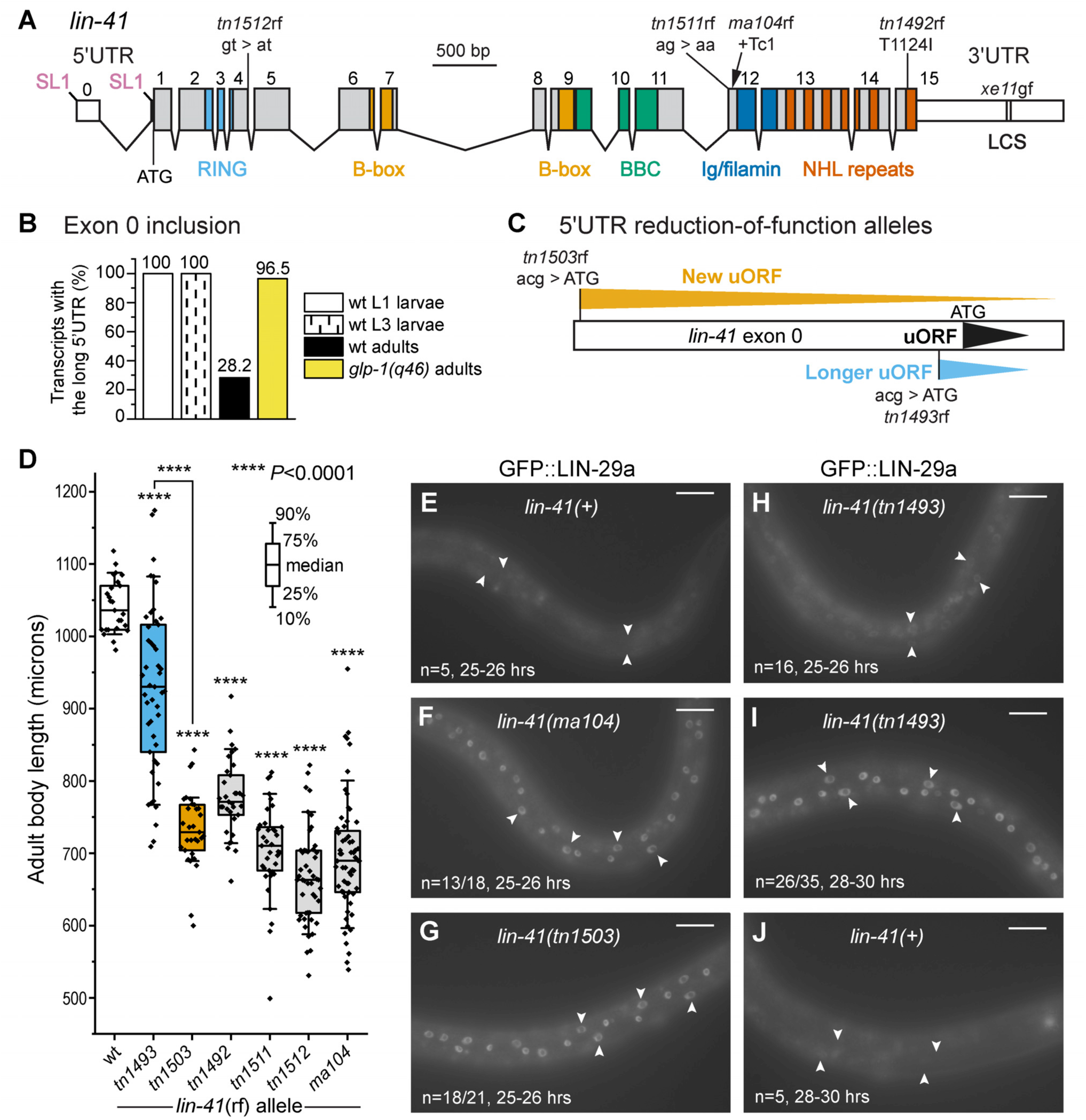
Novel *lin-41* reduction-of-function (rf) alleles map to a uORF-containing exon. (A) Gene structure of *lin-41*, showing LIN-41 protein domains and the position of addition of the SL1 trans-spliced leader sequence for the *lin-41* long and short transcripts. The positions of selected *lin-41(*rf) mutations that affect coding potential or splicing are shown (B) The percentage of *lin-41* long transcripts containing exon 0 (see Table S1). (C) Exon 0 *lin-41(*rf) mutations that were isolated as *let-7* suppressor mutations. The weaker mutation, *lin-41(tn1493)* lengthens the *lin-41* uORF from seven to 10 amino acids. The stronger *lin-41(tn1503)* mutation introduces a new ATG codon, creating a new uORF. (D) Adult body-size length measurements. Dots are data points, horizontal lines are the median, bars are the interquartile range, and whiskers encompass 80% of the data points. Statistical analyses utilized an ANOVA with a post-hoc Games-Howell test. Raw data and statistics are in Table S8. (E) *lin-41*(rf) mutations cause the precocious expression of GFP::LIN-29a. Animals were examined at the indicated times, post-hatch at 20°C. The number of animals scored is indicated. Arrowheads indicate example hypodermal nuclei. Bars, 20 μm.

As mentioned above, *lin-41* has soma-specific functions as part of the heterochronic gene regulatory pathway (Reinhart et al., 2000; Slack et al., 2000), as well as germline-specific functions during oogenesis, which commences at the L4-larval stage and continues into adulthood (Spike et al., 2014a,b; Tocchini et al., 2014). We found that wild-type L1 and L3-stage larvae exclusively expressed the *lin-41* long transcript containing exon 0 (Figs 2B and S1, Table S1). In contrast, wild-type adults expressed a majority (71.8%) of *lin-41* short transcripts and a minority (28.2%) of *lin-41* long transcripts (Figs 2B and S1, Table S1). Adults homozygous for the *glp-1(q46)* null mutation, which are sterile and produce only a few germ cells (Austin and Kimble, 1987), expressed the *lin-41* long transcript almost exclusively (96.5%; Figs 2B and S1, Table S1). Thus, the *lin-41* short transcript might be unique to the oogenic germline. *lin-41(tn1503*rf) and *lin-41(tn1493*rf) contain mutations in exon 0 and exclusively alter the 5’ UTR of the *lin-41* long transcript (Fig. 2C). *lin-41(tn1503)* is the stronger of the two mutations. *lin-41(tn1503*rf) animals, as well as other strong reduction-of-function *lin-41* alleles such as the canonical *lin-41(ma104)* allele (Slack et al., 2000), are shorter than wild-type adults and exhibit a pronounced Dumpy (Dpy) phenotype (Fig. 2D). In contrast, *lin-41(tn1493*rf) is milder with a weak and partially penetrant body-size defect (Fig. 2D). Remarkably, both *lin-41(tn1493*rf) and *lin-41(tn1503*rf) contained C to T mutations within exon 0 that introduced new initiator (ATG) codons. The mutation in *lin-41(tn1493*rf) is predicted to lengthen an endogenous upstream open reading frame (*lin-41* uORF) from seven to 10 amino acids (Fig. 2C). In contrast, *lin-41(tn1503*rf) is predicted to produce a new uORF (*lin-41* uORFn) of 51 amino acids (Fig. 2C). Several lines of evidence validated the assignment of *tn1493* and *tn1503* as *lin-41* alleles (see Materials and Methods). For example, they precociously expressed the *lin-41* target gene *lin-29a* in hypodermal epithelial cells at the L3 larval stage (Fig. 2E-J). GFP::LIN-29a is not expressed in hypodermal nuclei in the wild type during the L3 stage (Azzi et al., 2020) (Fig. 2E) but is expressed in hypodermal nuclei in most *lin-41(ma104*rf) larvae and *lin-41(tn1503*rf) larvae at this time (25-26 h post-hatch, Fig. 2E-G). Most *lin-41(tn1493)* L3 larvae also precociously express LIN-29a, but strong expression is observed slightly later (28-30 h post-hatch, Fig. 2H,I), at a time when wild type animals still failed to express LIN-29a in the hypodermis (Fig. 2J).

Studies in several experimental systems indicate that translation of uORFs can interfere with translational initiation at the downstream main open reading frame (Calvo et al., 2009; Zhong et al., 2024). To determine whether ribosomes are engaged within exon 0, we examined ribosome footprinting data from larval stages in the wild type and *let-7(n2853*ts) mutants (Hendriks et al., 2014; Aeschimann et al., 2017). We observed evidence for ribosome engagement within exon 0; however, ribosome occupancy was more pronounced within *lin-41* coding exons (Fig. S2; Table S2). The ribosome protected fragments observed within exon 0 mapped predominantly to regions 1 and 2 of exon 0 (see Fig. 3A). Ribosome protected fragments within exon 0 were also observed in *let-7(n2853*ts) mutants (Fig. S2; Table S2), suggesting that ribosome occupancy within exon 0 is independent of Let-7 regulation.

**Fig. 3.**
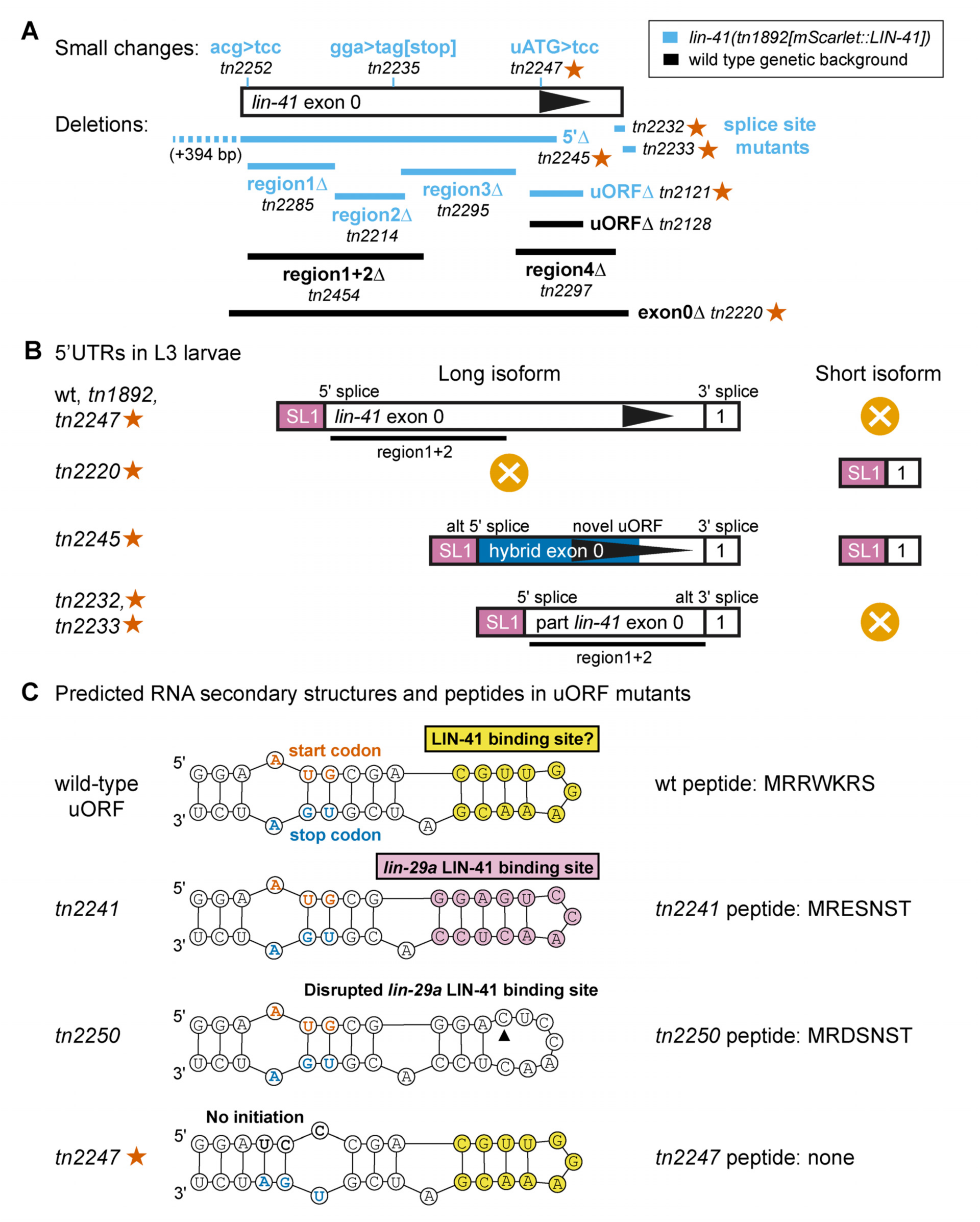
Genome editing to analyze functional elements within exon 0. (A) Genome edits in wild-type (black) and *lin-41(tn1892[mScarlet::lin-41])* (light blue) genetic backgrounds. Edits that gave rise to *lin-41(*gf) phenotypes are indicated with stars in all panels. (B) *lin-41* 5’UTRs found in the L3 larvae of specific *lin-41* mutants. Regions 1 and 2 are marked for reference (black bars). (C) Genome edits in the *lin-41(tn1892[mScarlet::lin-41])* genetic background used to assess potential roles of uORF peptide sequence, LIN-41 binding and uORF translation.

### Deletion of exon 0 causes a *lin-41* gain-of-function phenotype

The isolation of *lin-41* reduction-of-function mutations within exon 0 suggested that it might regulate *lin-41(+)* activity. We used genome editing to precisely delete exon 0 in an otherwise wild-type genetic background (Fig. 3A and Table S3) and found that animals homozygous for the *lin-41(tn2220[Exon 0Δ])* mutation exclusively expressed the *lin-41* short transcript at L3 and adult stages (Table S1, Fig. S1). *lin-41(tn2220[Exon 0Δ])* adult hermaphrodites exhibited an egg-laying defective (Egl) phenotype (Fig. 4A,E). At 20°C, 81% of *lin-41(tn2220[Exon 0Δ])* adults were Egl (n=59; Table S4). This Egl phenotype was semidominant (Fig. 4A); however, the penetrance of this semidominance was higher at 15°C (44% Egl at 15°C, n=50, Table S5). The *lin-41(tn2220[Exon 0Δ])* Egl phenotype resembled the phenotype caused by point mutations in two *let-7* control sequences [LCS, Ecsedi et al., 2015, *lin-41(xe11 [3’ LCS mut])*] in the *lin-41* 3’UTR (Fig. 4A, Table S4). These LCS mutations perturb downregulation of LIN-41 by Let-7. Consistent with a *lin-41* gain-of-function phenotype, *lin-41(tn2220[Exon 0Δ])* larvae failed to express GFP::LIN-29 in lateral epithelial cells at the L4 stage (Fig. 5A,B).

**Fig. 4.**
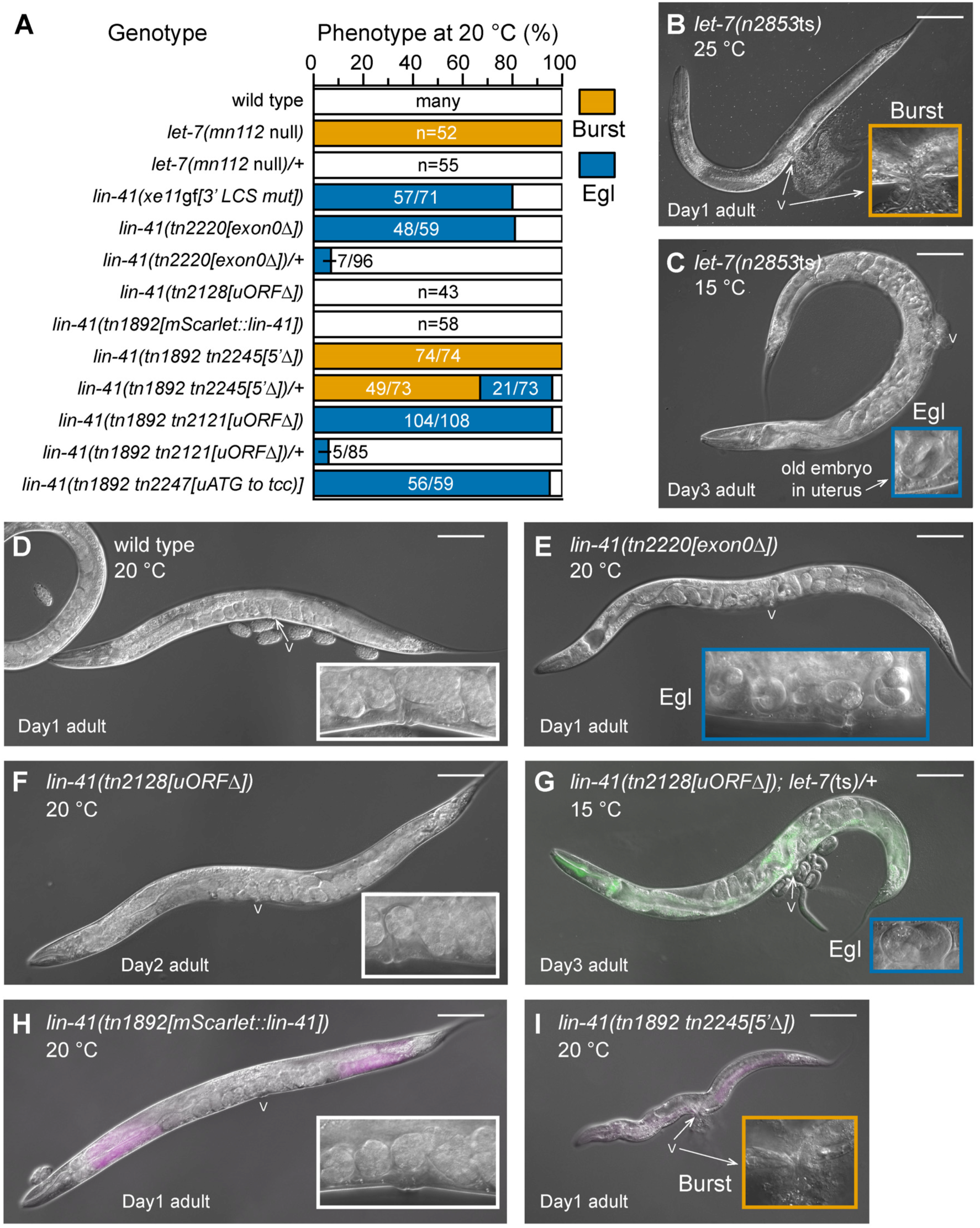
Phenotypes of *lin-41* exon 0 gain-of-function mutations and genetic interaction with *let-7*. (A) Phenotypes of the indicated *lin-41* mutations and comparison with the *let-7(mn112)* null allele and the *lin-41(xe11[3’ LCS mut])* gain-of-function allele that disrupts Let-7 recognition sequences in the *lin-41* 3’UTR. (B-I) DIC photomicrographs showing animals with *let-7(n2853*ts) reduction-of-function (B-C) and *lin-41(*gf) (E, G, I) phenotypes along with control strains (D, F, H) at the indicated temperatures and days of adulthood. Panels (G-I) are overlaid with fluorescence images showing GFP expression from the *tmC24* balancer for *let-7(n2853*ts) in (G) and mScarlet::LIN-41 expression in (H-I). Bars, 100 μm. Insets were magnified 3-fold to highlight egg-laying (C, E, G) and vulval bursting (B,I) phenotypes.

**Fig. 5.**
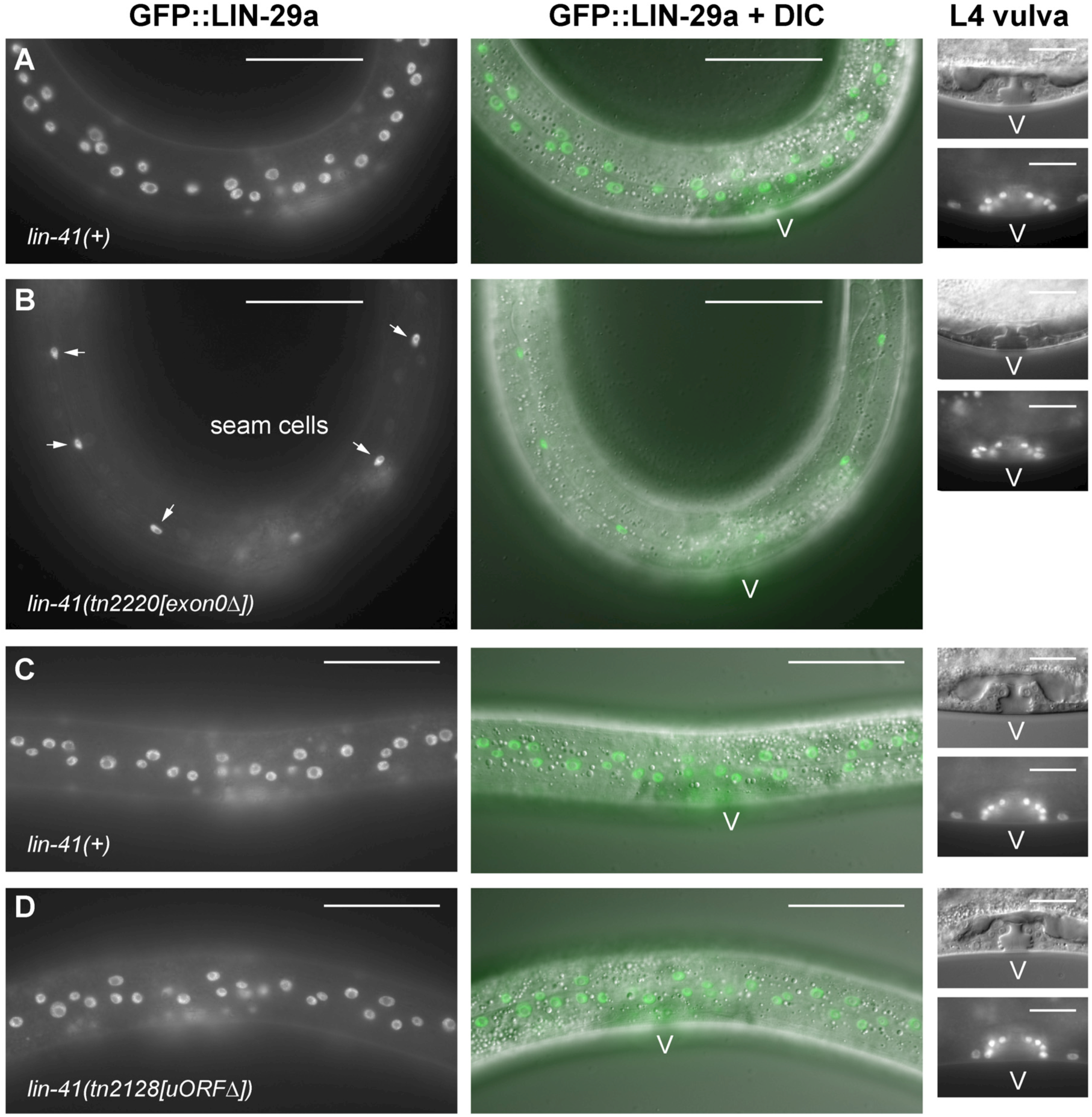
*lin-41(tn2220 [exon0Δ])* mutants fail to express GFP::LIN-29a in lateral epithelial cells. (A-D) GFP::LIN-29 expression at the L4 stage (left panels) in the indicated genetic backgrounds and merged images with DIC (middle panels). The panels on the right show DIC images (top) and LIN-29a::GFP expression (bottom) in the developing vulval region, highlighting the staging. Note, the *lin-41(tn2220 [exon0Δ])* mutation (B) does not affect GFP::LIN-29a expression in seam cells (arrows) or vulval cells (V). The *lin-41(tn2128[uORFD])* mutation does not affect GFP::LIN-29a expression. Bars, 50 mm for the left and middle panels; 20 mm for the vulval panels.

We observed that before *lin-41(tn2220[Exon 0Δ])* adults became Egl, they became lethargic, a behavioral phenotype associated with the process of molting (Lažetić and Fay, 2017). Because *let-7(n2853*ts) adults exhibit a supernumerary molt (Reinhart et al., 2000) and *lin-41(ma104*rf) L3 larvae exhibit precocious adult-stage molting (Slack et al., 2000), we examined the expression of *mlt-10p::gfp::pest*, which is expressed during the larval molts but not later during the adult stage (Frand et al., 2005). We found that a majority of *lin-41(tn2220[Exon 0Δ])* adults expressed *mlt-10p::gfp::pest* (72-73 h post-hatch at 22°C; 36 of 40 animals scored), whereas wild-type adults do not (Fig. S3A,C). We also used genome editing to delete the *lin-41* uORF (Fig. 3A). We observed no apparent defects in *lin-41(tn2128[uORFΔ])* hermaphrodites at any stage of development (Figs 4A,F, 5C,D and S3B, Tables S4 and S5). These observations suggest that deletion of exon 0 increases *lin-41(+)* activity but that the *lin-41* uORF is dispensable in an otherwise wild-type genetic background.

### Deletion of exon 0 and the *lin-41* uORF cause *lin-41* gain-of-function phenotypes in a sensitized genetic background

We sought to generate mutations within exon 0 in the context of fluorescently tagged *lin-41* alleles to monitor the effects of exon 0 mutations on LIN-41 expression. However, prior work showed that GFP fusions at the LIN-41 N-terminus cause gain-of-function phenotypes (Spike et al., 2018, 2022). For example, most *lin-41(tn1541[gfp::lin-41])* males exhibit a defect in tip retraction during the development of the male tail, producing a Leptoderan (Lep) tail (Spike et al., 2018). This defect is also found in gain-of-function *lin-41(*Lep) alleles affecting the N-terminal 39 amino acids of LIN-41 (Del Rio-Albrechtsen et al., 2006) and *lin-41(xe11 [3’UTR LCS mut])* males (Aeschimann et al., 2019). Animals that express mScarlet::LIN-41 had normal male tail development (Fig. S4) and hermaphrodite brood sizes comparable to the wild type (330 ± 33, n=18, versus 290 ± 25, n=19 for N2). We utilized *lin-41(tn1892[mScarlet::lin-41])* to examine the effects of exon 0 mutations on LIN-41 expression. Our analysis, described in detail below, demonstrated that *lin-41(tn1892[mScarlet::lin-41])* is a sensitized genetic background with elevated *lin-41(+)* activity relative to the wild type.

We attempted to generate a precise exon 0 deletion in the *lin-41(tn1892[mScarlet::lin-41])* but were unsuccessful after repeated attempts. However, we succeeded in isolating a deletion allele, *lin-41(tn1892[mScarlet::lin-41] tn2245[5’Δ])* that had a 543 bp deletion starting 394 bp upstream of exon 0, removing the SL1 splice acceptor sequence, and extending into exon 0 with the right breakpoint within the *lin-41* uORF (Fig. 3B and Table S3). *lin-41(tn1892[mScarlet::lin-41] tn2245[5’Δ])* L1-L3-stage larvae produce a novel *lin-41* long transcript (52.4% of *lin-41* transcripts), utilizing a new upstream SL1 splice acceptor sequence, as well as *lin-41* short transcripts (47.6% of *lin-41* transcripts), which are ordinarily never observed in L1-L3-stage larvae (Fig. 3B, Table S1, Figs S1 and S5). Remarkably, *lin-41(tn1892[mScarlet::lin-41] tn2245[5’Δ])* animals exhibited a semidominant lethal phenotype in which adults burst, phenocopying *let-7(mn112)* null mutants, with a difference being that *lin-41(tn1892[mScarlet::lin-41] tn2245[5’Δ])* is dominant but *let-7(mn112)* is recessive (Meneely and Herman, 1979; Fig. 4A,I and Table S4). Although 67% of *lin-41(tn1892[mScarlet::lin-41] tn2245[5’Δ])/+* heterozygotes burst and die early in adulthood (n=73, Table S4), a few progeny are produced. Interestingly, the *lin-41(tn1892[mScarlet::lin-41] tn2245[5’Δ])* long transcript contains a novel 18 amino acid uORF. We may have been unable to isolate a complete exon 0 deletion in the *lin-41(tn1892[mScarlet::lin-41])* background due to an even stronger dominant lethal phenotype.

Similarly, we used genome editing to delete the *lin-41* uORF in the *lin-41(tn1892[mScarlet::lin-41])* genetic background (Fig. 3A). We found that *lin-41(tn1892[mScarlet::lin-41] tn2121 [uORFΔ])* adult hermaphrodites exhibited a highly penetrant Egl phenotype that is semidominant with higher penetrance at 15°C (Fig. 4A and Tables S4 and S5). In addition, most *lin-41(tn1892[mScarlet::lin-41] tn2121[*uORFΔ]) males exhibited a Lep phenotype (93%, n=42), as compared to *lin-41(tn1892[mScarlet::lin-41])* males, none of which had Lep tails (n=33, Fig. S4). Most *lin-41(tn1892[mScarlet::lin-41] tn2121[uORFΔ])* adults also expressed the *mlt-10p::gfp::pest* molting marker in the adult stage (72-73 h, 51 of 57 animals examined, Fig. S3F), consistent with elevated *lin-41(+)* activity. We also generated a *lin-41* uORF mutation in the *lin-41(tn1892[mScarlet::lin-41])* genetic background in which the *lin-41* uORF ATG initiator codon is mutated to tcc (Fig. 3B), which is not considered to be an effective non-ATG initiator codon (Kearse and Wilusz, 2017). *lin-41(tn1892[mScarlet::lin-41] tn2247[uATG to tcc])* hermaphrodites exhibited an Egl phenotype (Table S4) and expressed the *mlt-10p::gfp::pest* molting marker in the adult stage (72-73 h, 42 of 45 animals examined, Fig. S3E). Taken together these results revealed that *lin-41(tn1892[mScarlet::lin-41])* constitutes a sensitized genetic background in which the *lin-41* uORF is required for normal development.

In addition to inhibiting translation initiation of the downstream main open reading frame, uORFs can also promote nonsense-mediated mRNA decay (NMD; Gaba et al., 2005). To examine whether the NMD pathway surveils *lin-41* transcripts via the *lin-41* uORF, we asked whether *smg-2(r908)*, a severe mutation in a gene essential for NMD (Page et al., 1999) mimics the effects of deleting the *lin-41* uORF or mutating the uORF start codon in the *lin-41(tn1892[mScarlet::lin-41])* genetic background. *smg-2(r908) lin-41(tn1892[mScarlet::lin-41])* adult hermaphrodites were egg-laying proficient; they did not have an Egl phenotype (n=60, Table S4). Consistent with this result, *smg-2(r908)* did not increase the expression of mScarlet::LIN-41 (Fig. S6A). Further, neither *lin-41(tn1503*rf) nor *lin-41(tn1493*rf) were suppressed by *smg-2(r908)* to a wild-type phenotype. These results suggested that the *lin-41* uORF can exert negative regulation upon *lin-41* in the absence of a functional NMD pathway. Additional results, described below, support this conclusion.

To examine the effect of the *lin-41* uORF on the regulation of *lin-41* activity, we used confocal and wide-field fluorescence microscopy to assess GFP::LIN-29a expression and mScarlet::LIN-41 fluorescence levels in L4-stage larvae, a key developmental stage at which *lin-41* is subject to regulation by the Let-7 microRNA (Reinhart et al., 2000). Relative to *lin-41(tn1892[mScarlet::lin-41])* control animals (Fig. 6A-B), *lin-41(tn1892[mScarlet::lin-41] tn2121[*uORFΔ]) L4 larvae showed a pronounced decrease in GFP::LIN-29a expression in the nuclei of lateral hypodermal epithelial cells (hyp7), although GFP::LIN-29a expression remained readily detectable in seam cell nuclei (Fig. 6C-D). Consistent with this observation, *lin-41(tn1892[mScarlet::lin-41] tn2121[uORFΔ])* animals appeared to exhibit elevated mScarlet::LIN-41 expression in hyp7 cells relative to *lin-41(tn1892[mScarlet::lin-41])* control animals, which had barely detectable levels of expression at the L4 stage (Fig. 6E-H). *let-7(n2853*ts) mutants exhibited an even greater apparent increase in mScarlet::LIN-41 expression, with detectable expression in both hyp7 and seam epithelial cells (Fig. 6I,J). We measured mScarlet::LIN-41 fluorescence in the heads of L4 animals to quantitatively determine the effects of various exon 0 mutations on *lin-41* expression (Figs 7, S6B and S7). Deletion of the *lin-41* uORF resulted in an average 2.4-fold increase in mScarlet::LIN-41 levels relative to control animals, with a similar 2.3-fold increase in *lin-41(tn1892[mScarlet::lin-41] tn2247[uATG to tcc])* mutants (Figs 7D-E and S6B). *let-7(n2853*ts) mutants displayed a 2.8-fold increase in mScarlet::LIN-41 levels (Figs 7F and S6B), whereas *lin-41(tn1892[mScarlet::lin-41] tn2121[*uORFΔ*]); let-7(n2853*ts) double mutants exhibited a significantly larger 4.2-fold increase (Figs 7G and S6B). Finally, the largest increase in mScarlet::LIN-41 levels, 5.4-fold relative to controls, was observed in *lin-41(tn1892[mScarlet::lin-41] tn2245[5’Δ])* mutants (Fig. S7), which lack most of exon 0 and exhibit a dominant bursting phenotype. Together, these results indicate that the *lin-41* uORF functions as a negative regulator of *lin-41* expression and activity and contributes to the regulatory function of exon 0.

**Fig. 6.**
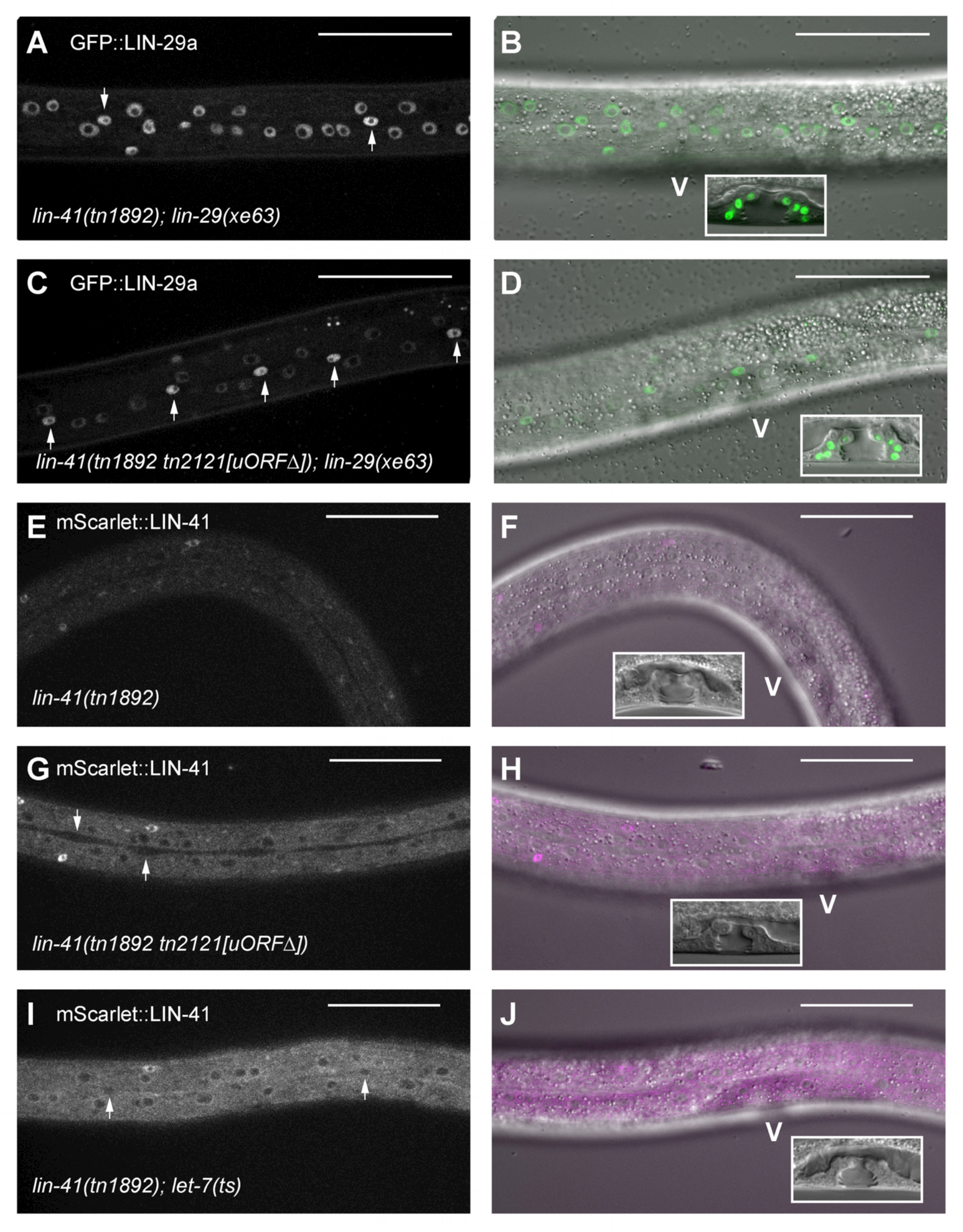
Deletion of the *lin-41* uORF increases expression of mScarlet::LIN-41 and reduces expression of GFP::LIN-29a. (A-D) GFP::LIN-29 expression in L4-stage larvae of the indicated genotypes (A,C) and shown merged with the DIC image (B,D). (E-J) mScarlet::LIN-41 expression in L4-stage larvae of the indicated genotypes (E,G,I) or shown merged with the DIC image (F,H,J). The boxed regions in the panels on the right show closeups of the vulval region (V), which are used in staging the animals. Arrows indicate seam cell nuclei. Note, the deletion of the *lin-41* uORF does not appear to affect the accumulation of GFP::LIN-29a in seam cell nuclei (C) or increase mScarlet::LIN-41 expression in seam cells (G). Bars, 50 μm.

**Fig. 7.**
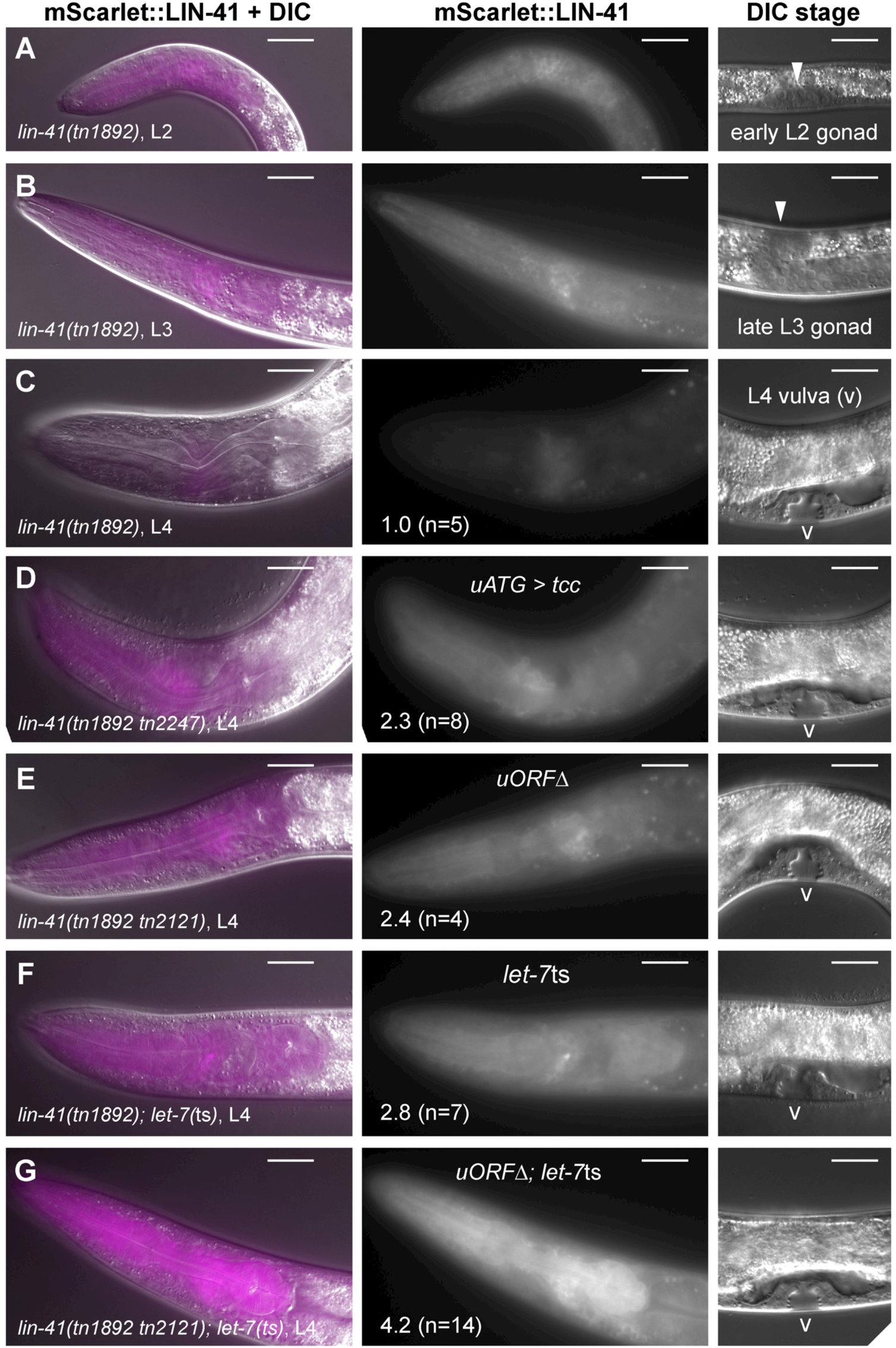
Effect of *lin-41* uORF mutations on mScarlet::LIN-41 fluorescence levels. (A-G) mScarlet::LIN-41 expression in the head regions of animals of the indicated genotypes and stages shown merged with DIC images (left panels) and in grayscale (middle panels). The panels at the right show images used in staging the animals. The stages examined were L2 (A), L3 (B) and L4 (C-G). Mean relative fluorescence values are indicated for the L4 stage (middle panels). Gonad (arrowheads) and vulva (V) are indicated. Bars, 20 μm.

### Exon 0 and *let-7* coordinately regulate *lin-41*

Exon 0 and *let-7* negatively regulate *lin-41*. We therefore examined whether deletions affecting exon 0 or the *lin-41* uORF enhance *let-7* mutant alleles (Fig. 8, Tables S4 and S5). Indeed, 52% of *lin-41(tn2128[uORFΔ]); let-7(n2853*ts*)/+* animals exhibited an Egl phenotype at 15°C, whereas *lin-41(tn2128[uORFΔ])* single mutants and *let-7(n2853*ts*)/+* heterozygotes developed normally (Figs 4G and 8A, Table S5). In addition, while only 10% of *let-7(n2853*ts) single mutants exhibited a Burst phenotype at 15°C, 100% of the *lin-41(tn2128[uORFΔ]); let-7(n2853*ts) double mutants exhibited this more severe phenotype (Table S5). We also found that 50% of *lin-41(tn2128[uORFΔ]); let-7(mn112)/+* hermaphrodites exhibited an Egl phenotype at 20°C even though the *let-7(mn112)* null allele is recessive (Fig. 8B and Table S4). Finally, we determined that, at 20°C, *lin-41(tn2220[Exon 0Δ])/+*; *let-7(n2853*ts)*/+* double heterozygotes exhibit a higher penetrance of the Egl phenotype (71%) than expected from the combined phenotypes of each heterozygote (Fig. 8B and Table S4). These genetic interactions provide an additional line of evidence that the 5’-regulatory exon, exon 0, plays a critical role needed for tight regulation of *lin-41* by Let-7.

**Fig. 8.**
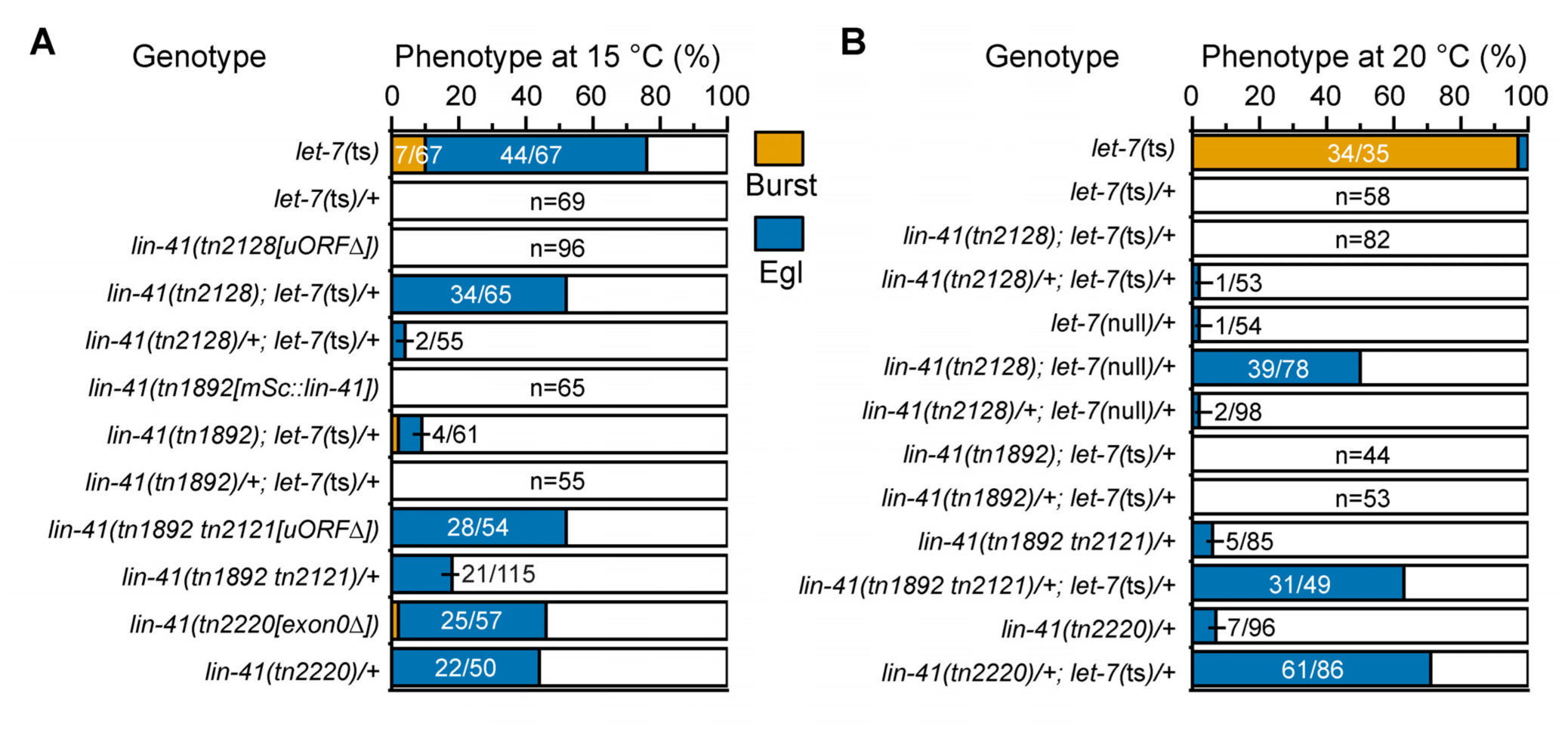
Genetic interactions between mutations affecting exon 0 and *let-7* mutations. Phenotypes of the indicated mutants and double mutant combinations at 15°C (A) and 20°C (B). Table S8 contains parent-of-origin and replicate information for all strains and genotypes that were analyzed.

### Intron 0 deletion imposes uORF-dependent negative regulation on *lin-41* during oogenesis

To test whether exon 0 could confer negative regulation on *lin-41* in the oogenic germline where the Let-7 microRNA is not expressed (Lau et al., 2001), we used genome editing to delete intron 0 so that all germline *lin-41* transcripts would include exon 0 (Fig. 9A). Whereas wild-type adults express a mixture of *lin-41* short and long transcripts in approximately a 4:1 ratio, *lin-41(tn2333[*intron 0Δ]) adults exclusively express *lin-41* long transcripts (Table S1, Fig. S1). There are two related effects of deleting intron 0 on oogenesis. First, *lin-41* transcript abundance in oocytes is decreased by ∼82% (Figs 9B and S8), possibly due to the removal of an E2F transcription factor binding-site located in the intron (Kudron et al., 2024). Second, the gonads produced small abnormal oocytes, and the animals exhibited a marked reduction in fertility (Figs 9C and S9A,B,E). When the *lin-41* uORF ATG is mutated to tcc in the context of the intron 1 deletion [e.g., *lin-41(tn2333[intron 0Δ] tn2353 [uATG to tcc])*], fertility is partially restored (Fig. 9C). We found comparable *lin-41* transcript levels in the oocytes of *lin-41(tn2333[intron 0Δ])* and *lin-41(tn2333[intron 0Δ] tn2353[uATG to tcc])* animals (Fig. 9B). Based on these observations, we conclude that the improved fertility of the double mutant is not due to an increase in *lin-41* transcript levels. This provides a second line of evidence that *lin-41* transcripts containing exon 0 and the uORF are not substrates for NMD, since comparable levels of *lin-41* transcripts are seen in oocytes when the *lin-41* uORF is present (*lin-41(tn2333[intron 0Δ])* mutant) and absent (*lin-41(tn2333[intron 0Δ] tn2353 [uATG to tcc])* mutant).

**Fig. 9.**
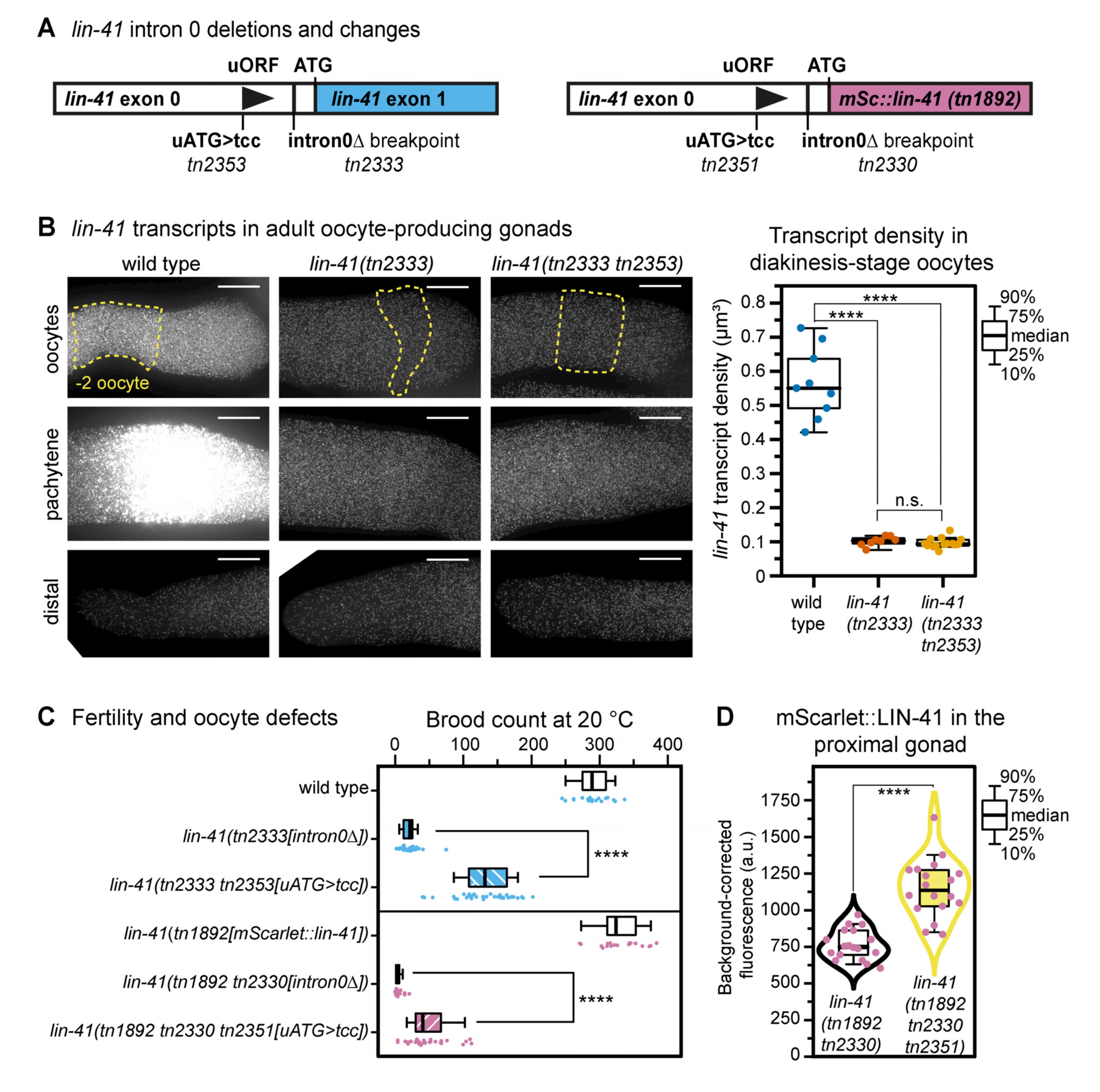
Intron 0 deletion causes uORF-dependent defects in oogenesis. (A) Intron 0 deletions made in wild-type (*tn2333*) and *lin-41(tn1892)* genetic backgrounds (*tn2330*). Mutations that change the *lin-41* uORF ATG to tcc are indicated (*tn2353* and *tn2351*). (B) *lin-41* mFISH in the indicated regions of the gonad in the indicated mutants (left panels), with quantification shown at the right. (C) Brood-size measurements in the indicated genotypes. (D) mScarlet::LIN-41 fluorescence measurements. For all graphs, dots are data points, boxplots with whiskers show the median, interquartile range and 80% of the data points. Statistical analyses employed an ANOVA with a post-hoc Games-Howell test (B, C) or a *t*-test with Welch’s correction (D), ****P<0.0001. Raw data and statistics are in Tables S7 and S8.

To examine the effects of deleting intron 0 on LIN-41 protein levels, we generated an intron 0 deletion in the *lin-41(tn1892[mScarlet::lin-41])* genetic background. mScarlet::LIN-41 was decreased by ∼84% in the proximal gonad of *lin-41(tn1892[mScarlet::lin-41] tn2330 [intron 0Δ])* adults relative to control animals (Fig. S9C), and these mutants exhibited very low fertility that was again improved by the *lin-41* uORF uATG to tcc mutation (Fig. 9C). We compared mScarlet::LIN-41 fluorescence intensity levels in the proximal gonads of both mutants and found that *lin-41(tn1892[mScarlet::lin-41] tn2330 [intron 0Δ] tn2351 [uATG to tcc])* germlines express approximately 1.5-fold more mScarlet::LIN-41 than *lin-41(tn1892[mScarlet::lin-41] tn2330 [intron 0Δ])* germlines (Figs 9D and S9E,F). Taken together, these results indicate that the *lin-41* uORF can impose negative regulation upon *lin-41* translation in the apparent absence of the Let-7 microRNA and without an effect on *lin-41* mRNA levels.

### Exon 0 contains evolutionarily conserved redundant elements

The analysis described above indicated that exon 0 negatively regulates *lin-41* activity and without this regulation, the Let-7 microRNA is unable to fully repress LIN-41 translation to enable normal development. To assess conservation, we analyzed the structure of the *lin-41* gene in Caenorhabditid nematodes that diverged from *C. elegans* approximately 80-100 million years ago (Stein et al., 2003; Hillier et al., 2007; Stevens et al., 2019). Exon 0 is highly conserved (Figs 10 and S10). We divided exon 0 into four regions; regions 1, 2 and 4 contain sequences that are highly conserved, whereas region 3 is divergent (Figs 3A, 10 and S10). Region 4 contains the *lin-41* uORF. All the species we examined contained a *lin-41* uORF, which varied from three amino acids to nine amino acids (Fig. 10B).

**Fig. 10.**
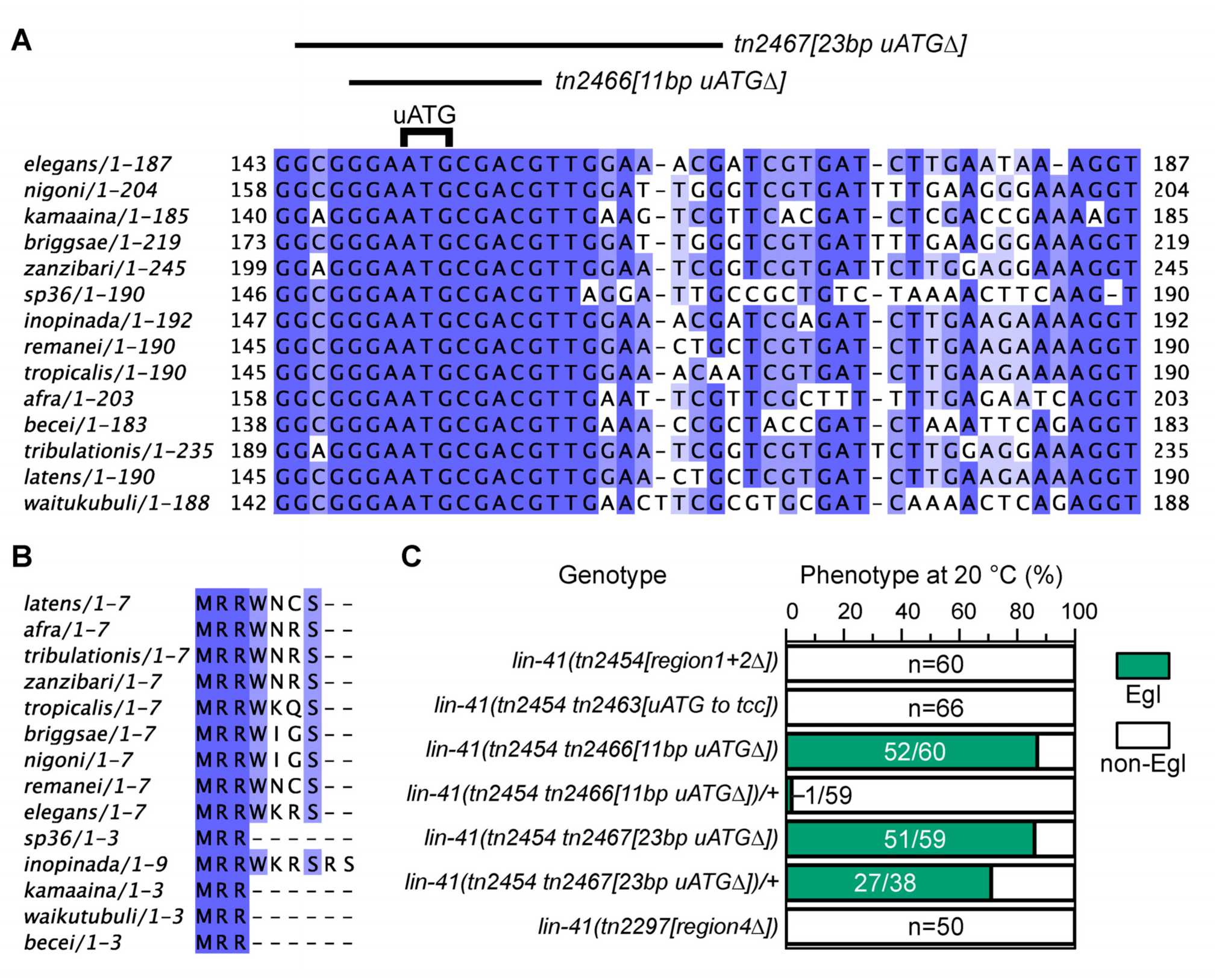
Exon 0 is conserved in Caenorhabditid nematodes and contains redundantly acting elements. (A) Sequence alignment of part of region 4 of exon 0 in the indicated Caenorhabditid nematodes. Deletions made in the *lin-41(tn2454[region 1,2*Δ]) genetic background and the *lin-41* uORF start codon are indicated. A full alignment of exon 0 is shown in Fig. S10. (B) Conservation of the amino acid sequence of the *lin-41* uORF. (C) Phenotypes of the indicated strains.

The observation that deletion of the *lin-41* uORF in a wild-type genetic background resulted in normal development, whereas deletion of exon 0 caused an Egl phenotype, suggested that exon 0 contains additional regulatory sequences in addition to the *lin-41* uORF. Thus, we used genome editing in the wild type and the *lin-41(tn1892[mScarlet::lin-41])* sensitized genetic background to dissect regulatory elements within exon 0. Region 1 contains the start site of the novel 51 amino acid LIN-41 uORF in the *lin-41(tn1503)* reduction-of-function allele (Figs 2C and 3A). Because ACG can function as an alternative initiator codon, albeit at reduced efficiency (Kearse and Wilusz, 2017), and we found evidence that ribosomes associate with this region of the 5’UTR (Fig. S2), we used the sensitized *lin-41(tn1892[mScarlet::lin-41])* genetic background to test whether the novel *lin-41* ACG-start uORF is a functional element within exon 0. We mutated the putative alternative ACG initiator codon to tcc, introduced a stop codon within region 2, and deleted the entirety of region 1, which all displayed normal development (Fig. 3A and Table S4). Based on these results, we conclude that *lin-41* uORFn is not a functional feature of exon 0 if it is translated at all.

To test whether regions 2 and 3 are required, we deleted them individually in the sensitized *lin-41(tn1892[mScarlet::lin-41])* genetic background and observed normal development (Fig. 3A and Table S4). Region 3 encodes a potential AU RNA stem loop with a ΔG value of –12.3 kcal/mol (at 37°C). Because energetically stable RNA stem loops can inhibit translation by obstructing ribosome progression (Kozak, 1986, 1989; Bao et al., 2020), we also deleted region 3 in the context of *lin-41* uORF deletion to generate *lin-41(tn2128[uORFΔ] tn2293[region3Δ])* to assess potential redundancy. However, *lin-41(tn2128[uORFΔ] tn2293[region3Δ])* mutants exhibited normal development, ruling out this possibility (Table S4). Since region 4 contains the *lin-41* uORF, which is required for normal development in the *lin-41(tn1892[mScarlet::lin-41])* genetic background, we deleted region 4 in a wild-type genetic background to generate *lin-41(tn2297[region 4Δ])*. Since a wild-type phenotype was observed, region 4 too appears to be dispensable (Fig. 3A and Table S4). This suggested that multiple exon 0 sequence elements function together to enable Let-7 microRNA regulation.

In the *lin-41(tn1892[mScarlet::lin-41])* genetic background, we isolated two mutations, *tn2232* and *tn2233*, which delete the splice donor sequence of intron 0 (Fig. 3A,B). The transcripts produced by these mutants utilize a cryptic splice donor sequence in the middle of exon 0 upstream of the *lin-41* uORF (Fig. 3B), such that only regions 1 and 2 are retained (Tables S1 and S3). Both *lin-41(tn1892[mScarlet::3xflag::lin-41] tn2232[intron0 splice donor* Δ1]) and *lin-41(tn1892[mScarlet::3xflag::lin-41] tn2233[intron0 splice donor* Δ2]) exhibited an Egl phenotype (Fig. 3B and Table S4). However, this phenotype is not as severe as caused by the dominant lethal *lin-41(tn1892[mScarlet::lin-41] tn2245[5’Δ])* mutant. This result suggested that conserved regions 1 and 2 are sufficient to mediate some repressive function by themselves. Yet deletion of regions 1 and 2 together in a wild-type background resulted in a wild-type phenotype, indicating that the rest of exon 0 is sufficient to mediate repression in that genetic background (Figs 3A, 10C and Table S4). To address redundancy, we made deletions that include a highly conserved block of nucleotide sequence and the *lin-41* uORF ATG in the absence of regions 1 and 2. Animals homozygous for *lin-41(tn2454 tn2466)* and *lin-41(tn2454 tn2467)* exhibited strong Egl phenotypes (Figs 10, S10 and Table S4). Interestingly, a *lin-41* uORF ATG to tcc mutation in the region 1–region 2 deletion resulted in a wild-type phenotype (Figs 3A, 10C and Table S4), suggesting the exon 0 contains at least three redundant functional elements including the *lin-41* uORF.

Close inspection of the *lin-41* uORF shows that it is embedded in an RNA sequence that has the potential to form a hairpin with a predicted LIN-41-binding site (Fig. 3C; Kumari et al., 2018). Thus, we addressed whether LIN-41 binding or autoregulation might potentially be involved in the translational regulation. To test this possibility, we first replaced the *lin-41* uORF sequences and the potential hairpin with an RNA hairpin from *lin-29a* (Fig. 3C), which binds LIN-41 (Aeschimann et al., 2017; Kumari et al., 2018). This genome editing replacement was conducted in the *lin-41(tn1892[mScarlet::lin-41])* sensitized genetic background to generate *lin-41(tn1892[mScarlet::lin-41] tn2241[lin-29a hairpin])*. In addition to changing the sequence of the potential RNA hairpin, this genome edit changed the protein coding sequence of the LIN-41 uORF from MRRWKRS to MRESNST. Since *lin-41(tn1892[mScarlet::lin-41] tn2241[lin-29a hairpin])* displayed a wild-type phenotype (Fig. 3C and Table S4), this experiment suggested that either the specific protein sequence of the LIN-41 uORF is not important or that LIN-41 binding is important. To distinguish between these possibilities, we replaced the *lin-41* uORF sequences and the potential hairpin with a mutant *lin-29a* RNA hairpin (Fig. 3C) that does not to bind LIN-41 (Kumari et al., 2018). This genome edit also changed the protein coding sequence of the LIN-41 uORF from MRRWKRS to MRDSNST. Since *lin-41(tn1892[mScarlet::lin-41] tn2250[lin-29a mutant hairpin])* displayed a wild-type phenotype (Fig. 3C and Table S4), we conclude that neither the potential LIN-41 binding nor the specific LIN-41 uORF protein sequence is required for Let-7 regulation of *lin-41* translation.

### Exon 0 regulates dauer formation

The regulation of GCN4 translation by amino acid starvation in budding yeast provides an archetypal example of uORF regulation (Dever et al., 2023). Thus, we asked whether exon 0 might regulate *lin-41* in response to nutrient availability. In *C. elegans*, starvation, high population densities and elevated temperatures cause the development of a stress-resistant L3-larval developmental state called dauer in which reproductive development arrests until conditions improve (Baugh and Hu, 2020). *lin-41(n2914)* null mutants form dauers inefficiently and the dauers they form exhibit morphological abnormalities (Cale and Karp, 2020; Wirick et al., 2021); however, post-dauer *lin-41(n2914)* adults are healthier than animals that have not transited through the dauer stage (Spike et al., 2014a). Based on these results, we hypothesized that *lin-41(+)* activity might promote the dauer developmental decision. Further, we imagined that exon 0 might repress *lin-41(+)* dauer-promoting activity under replete nutritional conditions. To test this possibility, we constructed *lin-41(tn2220[exon0Δ]); daf-7(e1372*ts) double mutants. *daf-7* encodes a TGF-β homolog that prevents dauer formation; *daf-7(e1372*ts) mutants exhibit a dauer-constitutive phenotype at 25°C (Riddle et al., 1981; Golden and Riddle, 1984; Ren et al., 1996). At 15°C on replete media with an abundant *E. co*li food source and low population density (∼5-20 worms/cm^2^), 13.7% of *daf-7(e1372*ts) mutants formed dauers (n=1987, Table 1). In contrast, under the same conditions, 36.3% of *lin-41(tn2220[*exon0Δ*]); daf-7(e1372*ts) double mutants formed dauers (n=787, Table 1, p<0.0001). The dauer-promoting activity of *lin-41(tn2220[*exon0Δ]) was dominant in that 33.7% of *lin-41(tn2220[exon0Δ])/+; daf-7(e1372*ts) animals formed dauers (n=1618, Table 1, p<0.0001). Neither *lin-41(tn2220[exon0Δ])* homozygotes nor *lin-41(tn2220[exon0Δ])/+* heterozygotes formed dauers (Table 1). We also investigated genetic interactions between *lin-41(tn2220[exon0Δ])* and *daf-2(e1370*ts). *daf-2* encodes an insulin/IGF-1 receptor homolog required for reproductive development (Riddle et al., 1981; Kimura et al., 1997). We constructed *lin-41(tn2220[exon0Δ]); daf-2(e1370*ts) double mutants and observed no enhancement of dauer formation at 15°C (Table 1). These results are consistent with the idea that *lin-41(+)* confers dauer-promoting activity that is ordinarily restrained by exon 0 under replete culture conditions.

**Table 1.** *lin-41(tn2220 [exon0Δ*gf]) enhances dauer formation at 15°C on replete medium.

| Genotype<sup>a</sup> | Dauer<sup>b</sup> (%) | N<sup>c</sup> |
| --- | --- | --- |
| *daf-7(1372)* | 13.7 | 1987 |
| *daf-7(e1372); lin-41(tn2220)/+* | 33.7<sup>d</sup> | 1618 |
| *daf-7(e1372); lin-41(tn2220)* | 36.3<sup>d</sup> | 787 |
| *daf-2(e1370)* | 0 | 2,080 |
| *daf-2(e1370); lin-41(tn2220)/+* | 0 | 1,339 |
| *daf-2(e1370); lin-41(tn2220)* | 0 | 719 |
| *lin-41(tn2220)/+* | 0 | 1,324 |
| *lin-41(tn2220)* | 0 | 717 |

<sup>a</sup>All genotypes contained *syIs601[ets-10p::gfp + ofm-1p::rfp]*. *ets-10p::gfp* is expressed in the intestine and neurons starting with the L2d-stage and continuing into the dauer stage. *ofm-1p::rfp* is expressed in coeloemocytes. *tmC18[tmIs1236(myo-2p::mCherry)]* was used as a balancer chromosome for *lin-* *41(tn2220)*.
<sup>b</sup>Animals were scored 96 h following egg laying at 15°C. Control animals had reached the L4-larval stage at this time.
<sup>c</sup>Compiled data from at least five biological replicates.
<sup>d</sup>*p*<0.00001 in comparison to the control, Chi-square test.

## DISCUSSION

Here, we used a genetic approach to investigate how Let-7 regulates *lin-41* in *C. elegans*. We identified a 5’-regulatory exon (exon 0) containing a uORF, as a key determinant of *lin-41* translational regulation in somatic cells. Our findings support a model in which exon 0 and its embedded uORF act with evolutionarily conserved redundant elements to limit translation initiation at the *lin-41* start codon, thereby permitting efficient Let-7-mediated repression of *lin-41* during the larval-to-adult transition. Consistent with this model, precise deletion of exon 0 caused egg-laying defects and inappropriate activation of the molting program in adults. In a sensitized genetic background, inactivation of the exon 0 regulatory module caused dominant-lethal “bursting,” phenocopying a *let-7* null mutation.

These findings reveal that the canonical Let-7 —| *lin-41* —| *lin-29* heterochronic pathway depends not only on Let-7-binding sites in the 3’UTR, but also on a 5’UTR-regulatory exon to achieve robust translational repression. To capture this behavior quantitatively, we modeled LIN-41 protein output with a rheostat-like relationship:

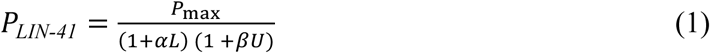

where *L* denotes Let-7 activity, *U* denotes exon 0 regulatory strength, P_max_ is maximal LIN-41 expression and *α* and *β* are coefficients specifying the strength of repressive inputs (Bintu et al., 2005). This formulation predicts that Let-7 activity and exon 0-mediated regulation combine multiplicatively to reduce LIN-41 expression. Using estimates derived from mScarlet::LIN-41 fluorescence measurements and the phenotypes of several key genetic backgrounds (see Materials and Methods), we generated a three-dimensional plot of LIN-41 expression as a function of Let-7 activity and exon 0 strength (Fig. S11 and Movie 1). The model predicts nonlinear graded reductions in LIN-41 levels when Let-7-mediated and exon-0-mediated repression operate simultaneously. Thus, exon 0 functions as a translational rheostat that attenuates translation initiation at the downstream *lin-41* start codon, enabling Let-7 to reduce *lin-41* expression during L4 to a range compatible with normal development.

Consistent with a role for exon 0 in tuning LIN-41 expression levels, we observed genetic interactions between *lin-41* exon 0 gain-of-function mutations and *let-7* mutant alleles. In sensitized genetic backgrounds, removal of the *lin-41* uORF increased LIN-41 protein abundance during the L4 stage, reduced LIN-29a accumulation and resulted in adult molting-related defects. Conversely, lengthening the *lin-41* uORF or introducing a novel uORF within exon 0 produced *lin-41* reduction-of-function phenotypes, including precocious LIN-29a expression and suppression of a *let-7* reduction-of-function mutation. Together with ribosome-footprinting data showing ribosome engagement within exon 0 (Aeschimann et al., 2017; Hendriks et al., 2014), these findings support a model in which translation of the *lin-41* uORF, together with conserved sequences within exon 0, limits reinitiation at the downstream *lin-41* start codon. One possibility is that Let-7–miRISC, recruited via the LCSs in the 3’UTR, acts on ribosome scanning or translation initiation to further reduce initiation from the downstream LIN-41 start codon. In this manner, the collaboration of 5’UTR and 3’UTR regulation would convert graded changes in uORF translational initiation to an on-off switch for the control of a dosage-sensitive developmental regulator.

Our analysis also shows that exon 0 can confer negative regulation on *lin-41* in other cell and development contexts that apparently do not involve the Let-7 microRNA, such as oogenesis and the dauer developmental decision. Deleting intron 0 forced all germline transcripts to include exon 0, which resulted in lower LIN-41 protein levels and severe oogenesis defects. Remarkably, these oogenesis defects were partially rescued by mutating the *lin-41* uORF start codon, which also increased LIN-41 protein levels. In the dauer-prone *daf-7(*ts) genetic background the deletion of exon 0 dominantly promoted dauer formation in *daf-7* mutants at 15°C under replete nutritional conditions. This result is consistent with the idea that *lin-41(+)* confers dauer-promoting activity (Cale and Karp, 2020; Wirick et al., 2021) that is normally restrained by exon 0 regulation under favorable growth conditions. Taken together these observations suggest that the exon 0–*lin-41* uORF module can be deployed independently of the Let-7 microRNA to adjust *lin-41* output to physiological and environmental cues. This regulatory principle might extend to LIN-41/Trim71-Let-7 circuitry in other species. uORFs are found in 40-50% of human 5’UTRs (Calvo et al., 2009). In fact, ribosome footprinting demonstrated that ribosomes are engaged on *LIN-41/Trim71* uORFs in embryonic stem (ES) cells (Popa et al., 2016). The Let-7 microRNA inhibits translation of LIN-41/Trim71 in ES cells and induced pluripotent stem cells and impacts differentiation (Worringer et al., 2014; Cuevas et al., 2015; Mitschka et al., 2015; Li et al., 2019). Further work will be needed to determine whether the regulatory principles we describe here in *C. elegans* for the regulation of *lin-41* translation by the Let-7 microRNA are more general.

## MATERIALS AND METHODS

### Strains and genetic analysis

#### Strain maintenance

The genotypes of the strains used in this study are reported in Table S6. *C. elegans* strain maintenance, strain constructions and genetic manipulations were conducted as described (Brenner, 1974). Routine strain maintenance and phenotypic analysis was generally conducted at 20°C except that temperature-sensitive mutant alleles were cultured at 15°C unless noted otherwise. CRISPR-Cas9 genome-editing manipulations were conducted at 22°C. Nematode growth medium (NGM) was composed of: 3 g l^-1^ NaCl, 20 g l^-1^ agar, 2.5 g l^-1^ peptone, 10 mg l^-1^ cholesterol, 6.25 mg l^-1^ nystatin, 0.2 g l^-1^ streptomycin, 1 mM CaCl_2_, 1 mM MgSO_4_, 12.5 mM KPO_4_ (pH 6.0). *E. coli* strain OP50-1 served as the food source. Genes and mutations are described in WormBase (Sternberg et al., 2024). In most cases, the balancer chromosome used to maintain *lin-41* mutant strains was *tmC18[dpy-5(tmIs1236[myo-2p::mCherry])]* (Dejima et al., 2018); however, *tmC18[dpy-5(tmIs1200[myo-2p::Venus])]* was used in crosses in which a green-fluorescent marker was needed. *let-7* mutant alleles were balanced by *tmC24[F23D12.4(tmIs1240[myo-2p::Venus])]* or *tmC24[F23D12.4(tmIs1233[myo-2p::mCherry])]*. Some strains and crosses used *tmC24* derivatives carrying the *unc-9(tm9718)* mutation, which could be rescued by *tmEx4950[unc-9(+) + vha-6p::gfp]*, enabling *tmC24*-carrying hemizygous males to mate. Strain constructions with *smg-2(r908)* utilized *unc-54(r293)*, which is suppressed when the NMD pathway is disrupted (Pulak and Anderson, 1993).

#### Brood size measurements and analysis of dauer formation

Brood-size measurements were conducted on the indicated genotypes to analyze fertility. L4-stage hermaphrodites were cultured individually and transferred to new media every 12-24 h. Viable progeny were counted before they reproduced. For analyzing dauer formation, approximately 20 gravid adult hermaphrodites of the indicated genotypes were cultured at 15°C on prechilled 5-cm diameter Petri dishes containing NGM medium and an *E. coli* food source at 15°C and allowed to lay eggs for 2 h after which the adults were removed. Dauer formation was scored 96 h after removing the parents. For analyzing the impact of the *lin-41(tn2220[exon 0Δ])* mutation on dauer formation, the *lin-41(tn2220[exon 0Δ]* parents used in the analysis were always heterozygous with *tmC18[dpy-5(tmIs1236[myo-2p::mCherry])]*. During these experiments we observed that *daf-7(e1372*ts), but not *daf-2(e1370*ts), suppressed the egg-laying defect of *lin-41(tn2220[exon 0Δ]* animals. Transit through the dauer stage has been observed to suppress heterochronic defects (Liu and Ambros, 1991), and passage through the dauer stage has been observed to ameliorate *lin-41* mutant phenotypes (Spike et al., 2014a).

#### Validation of lin-41(exon 0 rf) mutations as lin-41 alleles

The *lin-41(exon0* rf) mutations, *tn1493* and *tn1503*, which were isolated as dominant suppressors of *let-7(n2583*ts) (Spike et al., 2014a), map to chromosome I and contain single base substitutions in exon 0. Both *lin-41* exon 0 mutations fail to complement the *lin-41(n2914)* null mutation as shown by the following crosses. *lin-41(tn1503)/tmC18[tmIs1236]; tnEx265[str-1p::gfp]* and *lin-41(tn1493)/tmC18[tmIs1236]; tnEx265[str-1p::gfp]* males were crossed to *lin-41(n2914)/tmC18[tmIs1236]* hermaphrodites and F1 animals of genotypes *lin-41(n2914)/lin-41(tn1503); tnEx265[str-1p::gfp]* (n=18) and *lin-41(n2914)/lin-41(tn1493); tnEx265[str-1p::gfp]* (n=16) were examined and were observed to exhibit strong Dpy phenotypes. For both *lin-41* exon 0 reduction-of-function alleles, the heterozygotes with *lin-41(n2914)* were fertile; however, they were sicker for the stronger *tn1503* allele, which produced very few progeny.

#### Effect of the NMD pathway on exon 0 regulation of lin-41

Construction of *smg-2(r908) lin-41(tn1892) unc-54(r293)*: *lin-41(tn1892) / tmC18* males were crossed to *smg-1(r861) unc-54(r293)* hermaphrodites. *lin-41(tn1892) / smg-1(r861) unc-54(r293)* hermaphrodite cross progeny were allowed to reproduce by selfing. Progeny of genotype *lin-41(tn1892) unc-54(r293)* were isolated based on their Unc phenotype, their expression of mScarlet::LIN-41 and their failure to segregate non-Unc (*smg*-supressed) progeny. *lin-41(tn1892) unc-54(r293) / tmC18* males were then crossed to *smg-2(r908) unc-54(r293)* hermaphrodites. F1 Unc cross progeny of genotype *lin-41(tn1892) unc-54(r293) / smg-2(r908) unc-54(r293)* were allowed to reproduce by self-fertilization and suppressed F2 progeny of genotype *smg-2(r908) lin-41(tn1892) unc-54(r293*) were isolated and observed to have a wild-type phenotype. To assess suppression of *lin-41(tn1503)* and *lin-41(tn1493)* by *smg-2*, *lin-41(tn1503* or *tn1493) / tmC18* males were crossed to *smg-2(r908) lin-41(tn1892) unc-54(r293)* hermaphrodites. Hermaphrodites of genotype *lin-41(tn1503* or *tn1493) / smg-2(r908) lin-41(tn1892) unc-54(r293)* were allowed to reproduce by selfing. 30 Unc progeny of *lin-41(tn1503* or *tn1493) unc-54(r293) / smg-2(r908) lin-41(tn1892) unc-54(r293)* were cultured individually and suppressed animals of genotype *smg-2(r908) lin-41(tn1503* or *tn1493) unc-54(r293) / smg-2(r908) lin-41(tn1892) unc-54(r293)* were isolated. From these, *smg-2(r908) lin-41(tn1503 or tn1493) unc-54(r293)* homozygous progeny were isolated by virtue of their failure to express mScarlet::LIN-41. Neither *lin-41(tn1503)* nor *lin-41(tn1493)* was suppressed to wild-type by *smg-2(r908)*. *smg-2(r908) lin-41(tn1503) unc-54(r293)* hermaphrodites exhibited a strong Dpy phenotype; however, they appeared healthier than *lin-41(tn1503)* homozygotes. *smg-2(r908) lin-41(tn1493) unc-54(r293)* hermaphrodites exhibited an incompletely penetrant Dpy phenotype like *lin-41(tn1493)* hermaphrodites; however, their Dpy phenotype appeared mildly ameliorated.

#### Scoring lin-41(lf) body length and GFP::LIN-29a expression phenotypes

Body length was measured using animals selected at the L4 stage and cultured for 24 h at 20°C. These Day 1 adults were mounted in 0.1% levamisole on 2% agarose pads and imaged using a Nikon Ni-E microscope with a Plan Apo λ 10x/0.45 NA objective. Body length was measured in NIS-Elements (Nikon) by tracing a line along the midline of the animal from the tip of the nose to the base of the tail. Animals scored for GFP::LIN-29a expression were synchronized by collecting recently hatched larvae from manually picked embryos. GFP::LIN-29a expression in the hypodermis was imaged during the specified time windows (25-26 and 28-30 h post-hatching) using standardized imaging settings. Animals with “bright” GFP::LIN-29a fluorescence in hypodermal nuclei (mean intensity >1400 arbitrary units across 5 nuclei) were scored as GFP-positive. We also examined *lin-41(tn1503)* animals during an earlier time window (22-23 h post-hatching) and found that 20 of 30 animals were GFP-positive.

#### Scoring lin-41(gf) egg-laying and molting-related phenotypes

Egg-laying phenotypes were scored at multiple time points during the adult stage. At 20°C, we scored at approximately 24 h post L4, 40 h post L4 and 48 h post L4. At 15°C, we scored at approximately 48 h post L4 and 72 hr post L4. Animals were scored as having an egg-laying defect if at any time point they accumulated late-stage embryos in the uterus (2-fold stage or later) or if the embryos hatched internally. *mlt-10p::gfp* expression was examined in animals that had been synchronized by hatching in the absence of food. Animals were examined at multiple developmental time points after feeding starved L1s at 22°C — 47-48 h (L4), 50-51 h (L4-adult), 65-66 h (adult) and 72-73 h (adult). Strong GFP expression at 72-73 h was interpreted as the inappropriate execution of elements of a molting program at the adult stage. Consistent with this interpretation, the affected animals exhibited adult-specific lethargic and uncoordinated movement behaviors even in the absence of the *mlt-10p::gfp* reporter. This observation is important because it has been reported that the *mlt-10p::gfp* reporter we utilized (*mgIs49*) can generate a synthetic molting defect in a *myrf-1(mg412)* genetic background (Katic et al., 2026). In examining genetic interactions between *lin-41* exon 0 alleles and *let-7* mutant alleles, we focused on genetic combinations in which *let-7* was heterozygous because we suspected that some *let-7(n2853*ts) strains appeared to fortuitously harbor a recessive enhancer mapping to the region balanced by *tmC24*.

#### Sequencing of mutant alleles

PCR was conducted with primers listed in Table S3 and Sanger sequencing was performed (Azenta Genewiz). For whole genome sequencing of *lin-41(tn1485)*, genomic DNA was prepared using the QIAGEN DNeasy Blood and Tissue Kit. Illumina libraries of genomic DNA were prepared and paired-end (2×150) sequenced by Azenta GENEWIZ to approximately 60-200x coverage. Adapter and low-quality sequences were removed using Trim Galore! version 0.6.0 (Krueger, 2019) and mapped to the BSgenome.Celegans.UCSC.ce11 (1.4.2) genome using BWA mem (0.7.17-r1188). Aligned reads were sorted and quality filtered with Samtools (1.21). Duplicate reads were identified and marked with MarkDuplicates from Picard Toolkit (Broad Institute, 2019; version 2.18.16). The GATK Haplotype caller (4.1.2.0) was used to identify sequence variations with respect to the reference ce11 genome (Li and Durbin, 2009; Li et al., 2009; McKenna et al., 2010).

### Genome editing

Most genome-editing experiments were conducted by microinjecting plasmids encoding guide RNAs (gRNAs), the pDD162 Cas9-expressing plasmid (Dickinson et al., 2013) and a single-stranded oligonucleotide DNA repair template into the gonads of adult hermaphrodites. DNA and RNA oligonucleotides were purchased from IDT. Plasmids that express gRNAs under the control of the U6 promoter were generated by inserting sequence-specific oligonucleotides into the pRB1017 vector backbone as described (Arribere et al., 2013). The sequences of the gRNAs and the repair templates are listed in Table S3. Some repair templates introduced recognition sequences for restriction endonucleases (see Table S3), which aided in screening for genome edits. Most genome-editing experiments employed a *dpy-10* co-conversion strategy using the gRNA plasmid pJA58 and the AF-ZF-827 oligonucleotide repair template (Arribere et al., 2013). The experiments to generate the AU stem loop deletion and the intron 0 deletion utilized *myo-2p::Tdtomato* as the co-injected marker (4 ng/μl in the injection mix). For the genome-editing experiments that utilized the *dpy-10* co-conversion strategy, the injection mixes contained pJA58 (7.5 ng/μl), AF-ZF-827 (0.5 μM), gRNA-encoding plasmid(s) (25 ng/μl), *lin-41* repair template (0.5 μM) and pDD162 (7.5 ng/μl). Day 1 adult wild-type or *lin-41(tn1892)* hermaphrodites were injected and progeny in the F1 or F2 generations were screened by PCR using Q5 DNA polymerase, Taq polymerase, or Phusion polymerase (New England Biolabs), depending on whether the PCR products were analyzed following restriction enzyme digestion (the buffer for Q5 DNA polymerase is incompatible with many restriction enzymes). PCR primer sequences are listed in Table S3. Candidates containing the desired genome edits were backcrossed using *tmC18[dpy-5(tmIs1236)] / +* males. Genome edits were validated by PCR and Sanger sequencing (Azenta Genewiz). In most cases, at least two genome edits of a particular type were isolated.

The intron 0 deletions were generated using ribonucleoprotein injections. crRNAs and tracrRNA (Alt-R CRISPR-Cas9 tracrRNA, cat#1073190) were purchased from IDT. Cas9-NLS-6His was expressed in *E. coli* using pNM2973 (a kind gift of Michael Nonet; Fu et al., 2014) and purified as described (Paix et al., 2015). The injection mixes contained: intron 0 donor crRNA (4 pmol/μl), intron 0 acceptor crRNA (4 pmol/μl), *lin-41* repair oligonucleotide (500 nM), tracrRNA (16 pm/μl), Cas9 (1.2 μg/μl), pMyoTdTomato (4 ng/μl).

#### Attempts to generate an exon 0 deletion in the lin-41(tn1892) genetic background

In a single injection experiment using 39 wild-type animals (strain N2), we screened 64 adult F1 rollers, identified 9 candidate deletions, and recovered 4 independent *lin-41* exon 0 deletions, including *lin-41(tn2220)*. We attempted to identify exon 0 deletions by injecting 48 *lin-41(tn1892)* animals with the same injection mix. However, many of the *lin-41(tn1892)* F1 progeny had *lin-41(*gf) Egl or Burst phenotypes and there was a paucity of F1 rollers. After failing to identify deletion candidates by PCR among 72 adult roller F1s, we identified and recovered the intron 0 splice donor mutants, *lin-41(tn1892 tn2232)* and *lin-41(tn1892 tn2233)*, based on their Egl phenotypes. We also failed to identify *lin-41(tn1892)* animals with a precise exon 0 deletion after injecting animals heterozygous for *lin-41(tn1892)* and the null allele *lin-41(n2914)*. 86 adults of genotype *lin-41(tn1892) / lin-41(n2914); fog-2(oz40) / +* were injected with the same injection mix. 220 F1 rollers were screened by PCR and none contained the exon 0 deletion. *lin-41(tn1892 tn2245)* adults were fortuitously recovered in a later experiment to isolate the uATG to tcc mutation in the *lin-41(tn1892)* genetic background based on their dominant bursting phenotype and balanced using *tmC18[dpy-5(tmIs1236)]*.

### Analysis of *lin-41* transcripts by RT-PCR and exome sequencing

Approximately 50 animals of the desired stages and genotypes were collected in a 100 μl volume of DEPC-treated H_2_O in a 1.5 microcentrifuge tube, and 0.7 ml of TRizol LS (Life Technologies) was added. The sample was frozen in liquid nitrogen and stored at –80°C. Adults were analyzed approximately 16 h after the L4 stage at 20°C. For isolation of L1- and L3-stage larvae, embryos were collected by alkaline hypochlorite treatment (20% bleach and 0.5 N NaOH), washed in M9 buffer and allowed to hatch overnight in 5-cm diameter Petri dishes containing 8 ml of M9 solution. L1-stage larvae hatched in the absence of food were plated on NGM medium containing *E. coli* as a food source. The L1-stage larvae analyzed were fed for 3 h and the L3-stage larvae were fed for 30 h, after which they were washed free of bacteria with M9 buffer and harvested for RNA purification as above. For *lin-41(tn1892[mScarlet::lin-41] tn2245[5’Δ])* it was not possible to isolate embryos by alkaline hypochlorite treatment because of its dominant lethal phenotype. Thus, L1-stage to L3-stage larvae were handpicked, making sure that none contained the *tmC18[dpy-5(tmIs1236[myo-2p::mCherry])]* balancer chromosome.

Frozen worms in Trizol LS were thawed in a 37°C water bath for approximately 1 min, vortexed for 2 min and frozen in liquid nitrogen. After four freeze-thaw cycles, the RNA/TRizol mixture was transferred to a pre-spun (12,000g for 1 min at room temperature) Phase Lock Gel Heavy 2 ml centrifuge tube (5 Prime, cat#2302830). After addition of 140 μl of chloroform, the tube was shaken vigorously (not vortexed) for 15 s and then centrifuged at 12,000g for 10 min at 4°C. The aqueous phase (∼350 μl) was transferred to an RNase-free tube and mixed with 1 volume of freshly prepared 70% EtOH. The sample was transferred to an RNeasy MinElute spin column (Qiagen) placed in a 2 ml collection tube and allowed to incubate at room temperature for 2 min. The column was centrifuged at 10,000 rpm for 1 min at room temperature and the flow through was discarded. 350 μl of Buffer RW1 (Qiagen) was added to the spin column and centrifuged at 10,000 rpm for 1 min. The RNA bound to the filter was then treated with RQ1 RNase-free DNase (1 unit per μl final, Promega) in a volume of 80 μl at room temperature for 15 min. The column was washed with 350 μl of Buffer RW1, followed by 500 μl of Buffer RPE (Qiagen) and 500 μl of freshly prepared 80% EtOH. The RNA was then eluted with 19 μl of RNase-free H_2_O, which was then pipetted back onto the column for a second elution.

cDNA was prepared immediately by adding 1 μl of 10 mM dNTP mix (10 mM each dNTP) and 1 μl of 50 μM random primers (hexamers, Promega) to 7 μl of purified RNA in a total volume of 13 μl. The RNA-primer mix was heated at 65°C for 5 min and then placed on ice for at least 1 min. Reverse transcription (RT) reactions, including a control lacking reverse transcriptase, were carried out by adding 4 μl 5x SSIV Buffer, 1 μl 100 mM DTT, 1 μl RNaseOUT (Invitrogen) and 1 μl SuperScript IV Reverse Transcriptase (200 units/μl, Invitrogen). The RT reactions were incubated at 23°C for 10 min followed by 50°C for 10 min. The enzymes were then inactivated by incubating at 80°C for 10 min. PCR was conducted using 2.5 μl of 1:100 or 1:20 dilutions of the cDNA in 25 μl PCR reactions using Q5 DNA polymerase (New England Biolabs). PCR reactions used the SL1 primer and the bx37R primer for wild-type genetic backgrounds and mScDG2 primer for *lin-41(tn1892)* genetic backgrounds. Primer sequences are listed in Table S3. PCR reactions used 35 cycles with annealing temperatures of 64°C and extension temperatures of 72°C, with a 2 min extension time. PCR products were analyzed on 2% agarose gels with 0.5x TBE buffer. PCR products were never observed when SuperScript IV Reverse Transcriptase was omitted from the cDNA synthesis reaction.

For DNA sequencing, PCR products were purified using QiaPrep 2.0 spin columns (Qiagen) using buffers prepared according to the manufacturer’s instructions. Because somatic *lin-41* transcripts are apparently of low abundance in adults, 10 50 μl PCR reactions were conducted and pooled to get sufficient quantities of cDNA for exome sequencing from *glp-1(q46)* adults. Similarly, 20 50 μl PCR reactions were needed to get sufficient quantities of cDNA from *lin-41(tn1892 tn2245)* L1-L3 larvae. Purified DNA was quantified using Qubit fluorometric quantification. Amplicon-EZ exome sequencing (Azenta Genewiz) was conducted using 250 bp paired-end reads or 300 bp paired-end reads for *lin-41(tn1892 tn2232)* and *lin-41(tn1892 tn2233)*. Sequence reads were imported into R (4.4.0) using the ShortRead (1.62.0) package (Morgan et al., 2009). For PCR products less than 500 bp, overlapping forward and reverse reads were assembled using PEAR (0.9.11) (Zhang et al., 2014) and read in as a single DNA sequence. The Biostrings (2.72.1) package was used to determine the orientation of the sequence reads relative to the oligonucleotide primers used for the PCR. The frequency of each unique sequence was determined using the tables() function in the ShortRead package. This method enabled us to quantify the ratio of *lin-41* long transcripts to *lin-41* short transcripts based on the relative number of reads in the exome sequencing data. Sanger sequencing of the cDNA PCR fragments was also done for *lin-41(tn1892 tn2232)* and *lin-41(tn1892 tn2233)* mutants, and as for the exome sequencing, only a modified version of the *lin-41* long transcript was observed (see Tables S1 and S3). For *lin-41(tn2454 tn2466)* and *lin-41(tn2454 tn2467)* only Sanger sequencing of PCR fragments was performed and only the *lin-41* long transcript was observed (Table S3).

### Reanalysis of ribosome profiling data

Published ribosome profiling data (accession numbers GSE52910, GSE52864, GSE52905 and GSE80159) from the wild type, *let-7(n2853*ts), *lin-41(xe11)* and *lin-41(xe11); let-7(n2853*ts) mutants (Hendriks et al., 2014; Aeschimann et al., 2017) were reanalyzed to assess whether ribosome-protected fragments mapped to exon 0. Downloaded sequence files were trimmed with Trim Galore! (2.2.0) (Krueger et al., 2019) using settings for small RNA libraries and mapped to the WBcel235/ce11 genome using bwa mem (0.7.17) (Li and Durbin 2009). Mapped reads were sorted with samtools (1.21) (Danecek et al., 2021). Exon specific counts were calculated with Rsubread (2.18.0) (Liao et al., 2019) and visualized using the Gviz (1.48.0) package (Hahne and Ivanek, 2016) and code available at https://github.com/gearh006/uORF. To compare samples, the numbers of ribosome-protected fragments mapping to *lin-41* exons were normalized according to the total number of ribosome-protected fragments mapping to *lin-41* exons in individual experiments.

### Microscopy, smFISH and image analysis

Most microscope images were acquired on a Nikon Ni-E microscope with a Plan Apo λ 10x/0.45 NA objective (Fig. S3), a Plan Apo λ 20x/0.75 NA objective (Fig. 4B-I), a Plan Fluor 40x/1.3 NA oil objective (Fig. S7), a Plan Apo λ 60x/1.4 NA oil objective (Figs 2E-J, 4B-I insets, 5, 7, S8 and S9) or a Plan Apo λ 100x/1.45 NA oil objective (Fig. 9B and S4). Image acquisition used a SOLA light engine (Lumencor), an ORCA-Fusion C14440 digital camera (Hamamatsu) and NIS-Elements software (Nikon Inc.). Images in Fig. 6A-J were acquired on a Nikon Ti2 inverted confocal microscope with a Plan Fluor 40x/0.8 NA objective (Fig. 6A-D), or a Plan ApoIR 60x/1.27 NA objective (Fig. 6E-J) a motorized stage and resonant scanner using NIS-Elements (Nikon Inc.). Fluorescence was detected using a four-channel hybrid gallium arsenide phosphide and photomultiplier detector system (A1-DUG, Nikon), which combines two gallium arsenide phosphide photomultiplier tubes with two conventional multi-alkali photomultiplier tubes. We quantified the average fluorescence intensity within regions of interest in 2D images using NIS-Elements AR version 6.02.03 (Nikon inc.). Each measurement was independently background corrected using the average fluorescence intensity of a nearby area outside the body of the worm. Multiple images were analyzed per genotype, as specified in text, figures and supplemental tables. The L4-stage larval head was segmented to include as many cells anterior to the posterior pharyngeal bulb as possible, including the pharynx and nerve ring, while avoiding autofluorescent gut tissue (Figs 7, S6 and S7). The adult gonad was segmented using one of two methods: (1) Outlining the gonad to create a thick “U” shape that included the most proximal LIN-41-expressing oocyte, extended posteriorly around the loop, and ended adjacent to the starting position (as in Fig 9D). (2) Measuring the mean intensity along a 6-μm polyline positioned in the center of the gonad and drawn using the reference positions described in method 1 (as in Fig S9C). The overall change in intensity was then estimated by calculating mean intensity over a fixed 325-μm distance from the oocyte starting position (as in Fig. S9D). Statistical analyses were performed in Origin 2023b (version 10.0.5.157; OriginLab, Northampton, MA, USA) using its “two-sample *t*-test” function and integrated Python engine.

SmFISH on dissected gonads was conducted essentially as described (Spike et al., 2026). Gonads were dissected from adult hermaphrodites in less than 10 min in a glass depression well in PBT [RNase-free 1x PBS, pH 7.4 (Invitrogen) + 1% Tween (Thermo Fisher Scientific Inc.)] and fixed for 25 min in PBT + 3.7% formaldehyde (Electron Microscopy Sciences). Fixed gonads were washed with PBT and transferred to a 1.5 ml non-stick RNase-free microcentrifuge tube (Invitrogen). Once transferred to microfuge tubes, fixed gonads were pelleted for 2 min at 500 *g* prior to subsequent buffer changes. After a final wash with 1 ml PBT, the fixed gonads were stored in 0.5 ml 100% ice-cold methanol at –20°C overnight or up to several days. Gonads were then gradually rehydrated using PBT and equilibrated with 1 ml prepared Stellaris Wash Buffer A (SMF-WA1-60, Biosearch Technologies) for at least 5 min. All Stellaris buffers were prepared as specified by the manufacturer. Gonads were finally resuspended in 100 μl Stellaris RNA FISH Hybridization buffer (SMF-HB1-10, Biosearch Technologies) containing deionized formamide and 90 nM Cal Fluor 610-labeled *lin-41* probes (Table S7). Gonads were hybridized at 37°C with shaking for 1-2 days and then washed sequentially with 1 ml (1) Stellaris Wash Buffer A for 30-40 min, (2) Stellaris Wash Buffer A containing 2 μl 1 mg/ml DAPI for 30 min, and (3) Stellaris Wash Buffer B (SMF-WB1-20, Biosearch Technologies) for 5 min. Gonads were quickly washed twice with 1 ml PBT and incubated for 10 min with PBT containing 5 μg /ml 488-labeled wheat germ agglutinin (WGA) to mark oocyte boundaries. After two additional 1-ml washes with PBT, gonads were resuspended in NPG mounting medium [0.5% w/v N-propyl gallate, 50% glycerol, 20 mM Tris pH 8.0] and mounted for microscopy in a 1:1 volume ratio of sample and VectaShield (H-1000, Vector Laboratories). SmFISH images in Fig. 9B show deconvolved maximum-intensity projections of full-depth z-series collected at 0.25-μm intervals and optimized for transcript counting. These images were deconvolved using the fast deconvolution method of NIS-elements (Nikon inc.) with a custom noise level of 42 and are visually optimized to show individual transcripts when transcript density is low. Single focal plane images of the same gonads are shown in Fig. S9. The latter images were acquired using the large-image function of NIS-elements and visually optimized to illustrate the dynamic pattern of *lin-41* transcript accumulation in wild type gonads.

*lin-41* transcript count and density were determined using the 3D Measurement module in NIS-elements v6.02.03 (Nikon Inc.), essentially as described (Spike et al. 2026). Images were sharpened using a Gauss-Laplace transformation (setting 1.7). Individual smFISH transcripts were identified using the “clustered spots” option with an expected spot diameter of 0.35 μm and a local contrast threshold setting of 50. Regions of interest containing one or more diakinesis oocytes were manually segmented in 2D, converted to 3D binary objects, and used for spot quantification within the segmented volume. For transcript density estimates, z-slices lacking predicted *lin-41* transcripts were removed from the beginning or end of each z-stack. Analyses were further restricted to well-compressed gonads with a final z-stack thickness of 14-20 μm to minimize variability in volume measurements. Raw transcript-count data and statistical tests are in Table S7.

### Phylogenetic analysis

The *C. elegans* exon 0 sequence was used in a blastn search in May 2026 in the following *Caenorhabditis* species in NCBI (Camacho et al., 2009). The blastn parameters were adjusted to find less similar matches (max target sequences = 10, expect threshold = 1-10, word size = 16). The species searched were (species, genome assembly name, submitted GenBank assembly): *C. afra,* nxCaeAfra1.1, GCA_963570955.1; *C. becei*, ASM5094810v1, GCA_050948105.1; *C. briggsae,* CB4, GCA_000004555.3; *C. inopinata*, SP34_v7, GCA_003052745.1; *C. kamaaina*, nxCaeKama1.1, GCA_964211945.1; *C. latens* ASM225923v3, GCA_002259235.3; *C. nigoni*, ASM2792064v1, GCA_027920645.1; *C. remanei*, CRPX506, GCA_010183535.1; *C. sp. 36* PRJEB53466; CNIP, GCA_946814055.1; *C. tribulationis*; nxCaeTrib1.1, GCA_977925995.1; *C. tropicalis*m NIC58_v2, GCA_043792875.1; *C. waitukubuli*, nxCaeWait2.1, GCA_965219145.1; and *C. Zanzibari*, nxCaeZanz1.1, GCA_963966625.1.

The blastn alignment was used to identify the chromosome encoding exon 0, and the chromosomal sequence was downloaded from NCBI. In RStudio (2025.09.0+387), the Biostrings package in Bioconductor (Gentleman et al., 2004; Huber et al., 2015) was used to extract the genome sequence surrounding the aligned exon 0 sequence and to put the sequence in the correct orientation. The exon 0 sequences were aligned using Kalign (Lassmann, 2020; Madeira et al., 2024). These alignments were used to identify the likely splice junctions for exon 0. Ape (3.1.7) was used to find and translate the uORF sequences (Davis and Jorgensen, 2022), and they were aligned with Clustal Omega (Madeira et al., 2024). All alignments were visualized using JalView version 2.11.5 (Waterhouse et al., 2009), and the final figures were made using Adobe Illustrator 2026 (30.1.0).

### Computational model

To model LIN-41 activity as the combined activities of Let-7 and exon 0, we employed a simple rheostat-based model that is structurally equivalent to partition-function-based promoter-occupancy models in statistical-mechanical treatments of gene regulation (Bintu et al., 2005). Accordingly, we expressed LIN-41 levels (*P_LIN-41_*) using equation (1), see Discussion, where *L* is Let-7 activity, *U* is exon 0 strength, *P*_max_ is maximal LIN-41 expression and *α* and *β* are coefficients that set the strength of each effect. At the L4 larval stage, when LIN-41 is downregulated, *L*=*U*=1 and we estimated *P*_LIN-41_ to be 1. Estimates for the values of α and β were guided by measurements of mScarlet::LIN-41 expression levels (Figs 7, S6B and S7BC) and the phenotypes of *lin-41* alleles and genetic interactions between *lin-41* mutant alleles and *let-7* mutant alleles (Tables S4 and S5) to yield values of α=1 and β=1.4. In the germline, in which L=U=0, P_LIN-41_=P_max_=4.8. Figures were made using Matplotlib version 3.11 (Hunter, 2007).

### Artificial intelligence (AI) statement

AI (Claude and Perplexity) was used to assist in generating computer code for statistical analyses; all AI-assisted code and analytical outputs were independently reviewed and validated by the authors. AI was also used to generate search terms and identify potentially relevant literature, which was independently located and assessed using PubMed and other appropriate sources. All cited references were read and verified by the authors. During manuscript preparation, AI was used solely for grammar and language editing of author-written text. All authors reviewed and revised the AI-assisted output and accept full responsibility for the accuracy, originality and integrity of the final manuscript.

### Accessibility statement

We took steps to improve the accessibility of microscopy images and diagrams for readers with color vision deficiencies. Key fluorescence micrographs are presented in grayscale where appropriate. All graphs and most line art use a colorblind-accessible color palette (Wong, 2011).

## ACKNOWLEDGEMENTS

This manuscript is dedicated to our colleague Robert K. Herman whose laboratory isolated the first allele of *let-7*. We thank Erika Tsukamoto for technical assistance with whole genome sequencing and Tejiri Agbamu assistance with genome editing. We thank Todd Starich, Tatsuya Tsukamoto and Ann Rougvie for advice and suggestions during the course of this work. Some strains were provided by the Caenorhabditis Genetics Center, which is funded by National Institutes of Health Office of Research Infrastructure Programs (P40 OD010440). Research reported in this publication was supported by the National Institute of General Medical Sciences of the National Institutes of Health under awards R35GM144029, R01GM122787 and R01HD120295. The content is solely the responsibility of the authors and does not necessarily represent the official views of the National Institutes of Health.

## AUTHOR CONTRIBUTIONS

Conceptualization: C.A.S., D.G.; Methodology, formal analysis and investigation: C.A.S, M.D.G., K.M.W., M.E.Z., N.C. D.G.; Writing, review and editing: C.A.S, M.D.G., K.M.W., D.G.; Supervision, project administration and funding acquisition: D.G.

## FUNDING

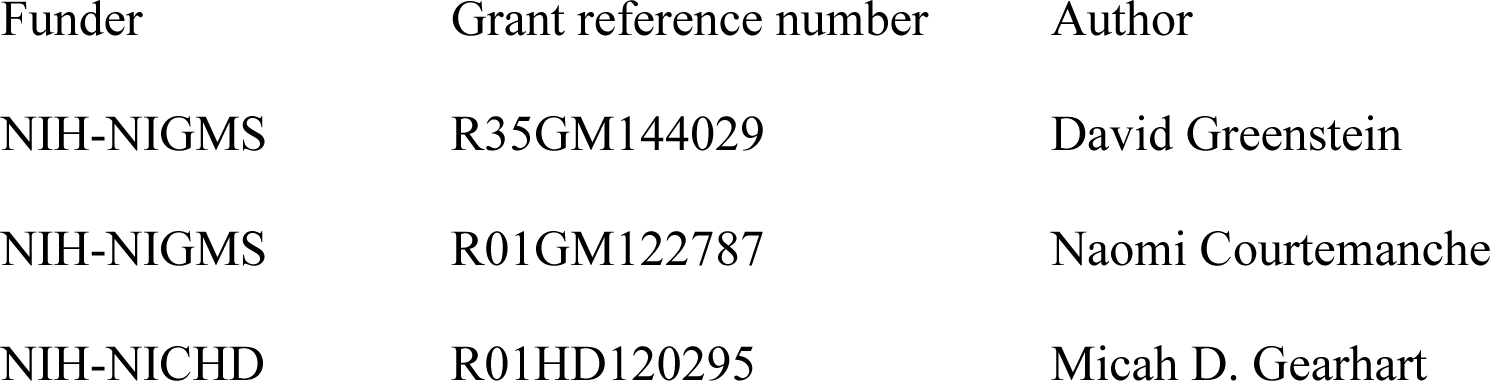

## DATA AND RESOURCE AVAILABILITY

Raw phenotypic data and associated statistical analyses are in Table S8. All strains are available from the authors or the Caenorhabditis Genetics Center upon request.

## Funding

### Funding Group

- Award Group:

- Funder(s): National Institutes of Health
- Award ID: R35GM144029
- Award ID: R01GM122787
- Award ID: R01HD120295

