## Supplemental Information for "Translation of a small upstream open reading frame functions as a rheostat for the regulation of *lin-41* by the Let-7 microRNA in *Caenorhabditis elegans*"

Contents:

8 Supplemental Tables

11 Supplemental Figures

1 Supplemental Movie

**Table S1** (related to Fig. 2). **The ratio of *lin-41* long and *lin-41* short transcripts in the wild-type and mutants**

| Genotype | Stage | *lin-41* long mRNA reads[a] | *lin-41* short mRNA reads[a] |
|---|---|---|---|
| WT (strain N2) | Adult | 112,612 (28.2%)[b] | 286,872 (71.8%)[b] |
| WT (strain N2) | L1 | 96,446 (100%)[c] | 0 (0%)[c] |
| WT (strain N2) | L3 | 115,709 (100%)[c] | 0 (0%)[c] |
| *glp-1(q46)* | Adult | 43,442 (96.5%)[c] | 1,591 (3.5%)[c] |
| *lin-41(tn2220 [Exon 0Δ])* | Adult | 0 (0%)[c] | 172,695 (100%) |
| *lin-41(tn2220 [Exon 0Δ])* | L3 | 0 (0%)[c] | 373,849 (100%)[c] |
| *lin-41(tn2333 [Intron 0Δ])* | Adult | 44,774 (100%)[b] | 0 (0%)[b] |
| *lin-41(tn2333 Intron 0Δ tn2353 uATG to tcc)* | Adult | 104,677 (100%)[c] | 0 (0%)[c] |
| *lin-41(tn1892[mScarlet::lin-41])* | Adult | 193,720 (34.6%)[b] | 365,668 (65.4%)[b] |
| *lin-41(tn1892[mScarlet::lin-41])* | L1 | 306,636 (100%)[c] | 0 (0%)[c] |
| *lin-41(tn1892[mScarlet::lin-41])* | L3 | 371,032 (100%)[c] | 0 (0%)[c] |
| *lin-41(tn1892[mScarlet::lin-41] tn2247 [uATG to tcc])* | Adult | 101,894 (40.6%)[c] | 149,312 (59.4%)[c] |
| *lin-41(tn1892[mScarlet::lin-41] tn2247 [uATG to tcc])* | L1 | 348,364 (100%)[c] | 0 (0%)[c] |
| *lin-41(tn1892[mScarlet::lin-41] tn2247 [uATG to tcc])* | L3 | 325,286 (100%)[c] | 0 (0%)[c] |
| *lin-41(tn1892[mScarlet::lin-41] tn2245 [5'Δ])* | L1-L3 | 8,115[c,d] (52.4%) | 7,361 (47.6%)[c,d] |
| *lin-41(tn1892[mScarlet::lin-41] tn2232)* | L3 | 53,611[c,e] (100%) | 0 (0%)[c,e] |
| *lin-41(tn1892[mScarlet::lin-41] tn2233)* | L3 | 60,024[c,e] (100%) | 0 (0%)[c,e] |

[a]Sequencing reads specific to *lin-41* long and *lin-41* short transcripts from exome sequencing of RT-PCR products.

[b]Aggregate data from two biological replicates.

[c]Data from a single biological replicate.

[d]As described in the text, *lin-41(tn1892[mScarlet::lin-41] tn2245 5'Δ)* is a 5'-deletion of exon 0, that removes the normal SL1 splice acceptor. SL1 is transpliced to a new upstream SL1 splice acceptor sequence. The right breakpoint is within the *lin-41* uORF sequence. Because of the dominant lethality of this mutant, we were unable to synchronize larvae using alkaline hypochlorite treatment. Consequently, a mixture of L1–L3-stage larvae was analyzed. The low number of reads from these samples is due to non-specific PCR products that do not originate from the *lin-41* gene.

[e]*lin-41(tn1892[mScarlet::lin-41] tn2232)* and *lin-41(tn1892[mScarlet::lin-41] tn2233)* delete the splice donor sequence of intron 0. Sequencing of the cDNA products indicates that a cryptic splice

donor sequence is used within exon 0, resulting in the removal of the *lin-41* uORF and retention of region 1 and region 2 in the transcript. No *lin-41* short transcripts are produced.

**Table S2** (related to Figs S2). **Reanalysis of ribosome profiling data.**

**Table S3** (related to Fig. 3). **Genome editing methods used to dissect the regulatory function of exon 0.**

**Table S4** (related to Fig. 4). ***lin-41*** **gain-of-function phenotypes and genetic interactions with *let-7* at 20º C.**

| Genotype [a] | Egl [b] | Burst [c] | Number Scored |
|---|---|---|---|
| **Homozygous mutant genotypes** | | | |
| 1. *let-7(n2853*ts*)* [d] | 3% | 97% [e] | n=35 |
| 2. *unc-3(e151) let-7(mn112)* | 0% | 100% | n=52 |
| 3. *lin-41(xe11[3'lcs1&2, c>t])* | 80% | 0% | n=71 |
| 4. *lin-41(tn1892[mScarlet::lin-41])* | 0% | 0% | n=58 |
| 5. *smg-2(r908) lin-41(tn1892) unc-54(r293)* | 0% | 0% | n=60 |
| 6. *lin-41(tn1892 tn2121[uORFΔ])* | 96% | 0% | n=108 |
| 7. *lin-41(tn1892 tn2214[exon0 region2Δ])* | 0% | 0% | n=48 |
| 8. *lin-41(tn1892 tn2232[intron0 splice donorΔ])* | 93% | 0% | n=58 |
| 9. *lin-41(tn1892 tn2233[intron0 splice donorΔ])* | 82% | 0% | n=65 |
| 10. *lin-41(tn1892 tn2235[exon0 stop codon])* | 0% | 0% | n=48 |
| 11. *lin-41(tn1892 tn2241[uORF>lin-29a stem loop1])* | 0% | 0% | n=50 |
| 12. *lin-41(tn1892 tn2245[5'Δ])* | 0% | 100% | n=74 |
| 13. *lin-41(tn1892 tn2247[uATG>tcc])* | 95% | 0% | n=59 |
| 14. *lin-41(tn1892 tn2250[uORF>mutant lin-29a stem loop1])* | 0% | 0% | n=50 |
| 15. *lin-41(tn1892 tn2252[exon0 acg>tcc])* | 0% | 0% | n=52 |
| 16. *lin-41(tn1892 tn2285[exon0 region1Δ])* | 0% | 0% | n=54 |
| 17. *lin-41(tn2128[uORFΔ]) tn2293[exon0 region3Δ])* | 0% | 0% | n=58 |
| 18. *lin-41(tn1892 tn2295[exon0 region3Δ])* | 0% | 0% | n=60 |
| 19. *lin-41(tn2128[uORFΔ])* | 0% | 0% | n=43 |

| | | | |
|---|---|---|---|
| 20. *lin-41(tn2220[exon0Δ])* | 81% | 0% | n=59 |
| 21. *lin-41(tn2297[exon0 region4Δ])* | 0% | 0% | n=50 |
| 22. *lin-41(tn2454[exon0 region1+2Δ])* | 0% | 0% | n=60 |
| 23. *lin-41(tn2454 tn2463[uATG>tcc])* | 0% | 0% | n=66 |
| 24. *lin-41(tn2454 tn2466[11bp uATGΔ])* | 87% | 0% | n=60 |
| 25. *lin-41(tn2454 tn2467[23bp uATGΔ])* | 86% | 0% | n=59 |
| ***lin-41/*+ genotypes** | | | |
| 26. *lin-41(tn1892 tn2121[uORFΔ])/tmC18* | 6% | 0% | n=85 |
| 27. *lin-41(tn1892 tn2245[5'Δ])/tmC18* | 29% | 67% [f] | n=73 |
| 28. *lin-41(tn2220[exon0Δ])/tmC18* | 7% | 0% | n=96 |
| 29. *lin-41(tn2454 tn2466[11bp uATGΔ])/tmC18* | 2% | 0% | n=59 |
| 30. *lin-41(tn2454 tn2467[23bp uATGΔ])/tmC18* | 71% | 0% | n=38 |
| ***lin-41; let-7ts/*+ genotypes** | | | |
| 31. *let-7(n2853*ts*)/tmC24* [d] | 0% | 0% | n=58 |
| 32. *lin-41(tn1892); let-7(n2853*ts*)/tmC24* | 0% | 0% | n=44 |
| 33. *lin-41(tn2128)*; *let-7(n2853*ts*)/tmC24* | 0% | 0% | n=82 |
| ***lin-41/+; let-7ts/*+ genotypes** | | | |
| 34. *lin-41(+)/tmC18*; *let-7(n2853*ts*)/tmC24* [g] | 0% | 4% | n=45 |
| 35. *lin-41(tn1892)/tmC18*; *let-7(n2853*ts*)/tmC24* | 0% | 0% | n=53 |
| 36. *lin-41(tn1892 tn2121)/tmC18; let-7(n2853*ts*)/tmC24* | 63% | 0% | n=49 |
| 37. *lin-41(tn2128)/tmC18*; *let-7(n2853*ts*)/tmC24* | 2% | 0% | n=53 |
| 38. *lin-41(tn2220)/tmC18*; *let-7(n2853*ts*)/tmC24* | 71% | 0% | n=86 |
| ***let-7(null)/*+ genotypes** | | | |

| | | | |
|---|---|---|---|
| 39. *lin-41(+); unc-3(e151) let-7(mn112)/tmC24* | 0% | 0% | n=55 |
| 40. *lin-41(tn2128); unc-3(e151) let-7(mn112)/tmC24* | 50% | 0% | n=78 |
| 41. *lin-41(+)/tmC18; unc-3(e151) let-7(mn112)/tmC24* | 2% | 0% | n=54 |
| 42. *lin-41(tn2128)/tmC18; unc-3(e151) let-7(mn112)/tmC24* | 2% | 0% | n=98 |

[a] Animal genotypes are abbreviated for clarity. Balancer chromosome *tmC24* is *let-7(+)* and refers to *tmC24[F23D12.4(tmIs1240) unc-9(tm9719)]*. Balancer chromosome *tmC18* is *lin-41(+)* and refers to *tmC18[dpy-5(tmIs1236)]*.

[b] Egg laying defective or "Egl" animals carried old embryos in their uterus 24-48 hours after being selected as L4-stage larvae. Some animals died and were consumed by the larval progeny that hatched inside the uterus. Other animals laid the embryos before they hatched. Both types were scored as "Egl".

[c] "Burst" animals leaked material through the vulva as young adults and died before making many embryos.

[d] Animals from parent strain DG5827 *let-7(n2853*ts*)/tmC24*.

[e] 26% of the "Burst" animals produced some progeny.

[f] 90% of the "Burst" animals produced some progeny.

[g] Animals from parent strain DG6062 *lin-41(+)/tmC18; let-7(n2853*ts*)/tmC24*.

**Table S5** (related to Fig. 4). ***lin-41* gain-of-function mutant phenotypes and genetic interactions with *let-7* at 15 °C.**

| Genotype [a] - strain of origin [b] | Egl [c] | Burst [d] | Number Scored |
|---|---|---|---|
| **Homozygous mutant genotypes** | | | |
| 1. *let-7(n2853*ts*)* - DG5827 | 66% | 10% | n=67 |
| 2. *let-7(n2853*ts*)* - DG6062 | 12% | 85% | n=33 |
| 3. *lin-41(tn1892[mScarlet::LIN-41])* | 0% | 0% | n=65 |
| 4. *lin-41(tn1892); let-7(n2853*ts*)* | 0% | 100% | n=58 |
| 5. *lin-41(tn1892 tn2121[uORFΔ])* | 52% | 0% | n=54 |
| 6. *lin-41(tn1892 tn2121)*; *let-7(n2853*ts*)* | 0% | 100% | n=12 |
| 7. *lin-41(tn2128[uORFΔ])* | 0% | 0% | n=96 |
| 8. *lin-41(tn2128)*; *let-7(n2853*ts*)* | 0% | 100% | n=47 |
| 9. *lin-41(tn2220[exon0Δ])* | 44% | 2% | n=57 |
| 10. *lin-41(tn2220); let-7(n2853*ts*)* | 0% | 100% | n=15 |
| 11. *lin-41(tn2454[region1+2Δ]* | 0% | 0% | n=57 |
| ***lin-41/+*** **genotypes** | | | |
| 12. *lin-41(tn1892 tn2121)/tmC18* | 18% [e] | 0% | n=115 |
| 13. *lin-41(tn2220)/tmC18* | 44% | 0% | n=50 |
| ***lin-41; let-7ts/+*** **genotypes** | | | |
| 14. *let-7(n2853*ts*)/tmC24* - DG5827 | 0% | 0% | n=69 |
| 15. *let-7(n2853*ts*)/tmC24* - DG6062 | 0% | 0% | n=30 |
| 16. *lin-41(tn1892); let-7(n2853*ts*)/tmC24* | 7% | 2% | n=61 |
| 17. *lin-41(tn1892 tn2121)*; *let-7(n2853*ts*)/tmC24* | 36% | 5% | n=22 |

| | | | |
|---|---|---|---|
| 18. *lin-41(tn2128)*; *let-7(n2853*ts*)/tmC24* | 52% | 0% | n=65 |
| 19. *lin-41(tn2220); let-7(n2853*ts*)/tmC24* | 52% | 0% | n=27 |
| ***lin-41/+; let-7ts/+* genotypes** | | | |
| 20. *lin-41(+)/tmC18; let-7(n2853*ts*)/tmC24* – DG6062 | 0% | 0% | n=30 |
| 21. *lin-41(tn1892)/tmC18; let-7(n2853*ts*)/tmC24* | 0% | 0% | n=55 |
| 22. *lin-41(tn1892 tn2121)/tmC18*; *let-7(n2853*ts*)/tmC24* | 65% | 0% | n=22 |
| 23. *lin-41(tn2128)/tmC18; let-7(n2853*ts*)/tmC24* | 4% | 0% | n=55 |
| 24. *lin-41(tn2220)/tmC18; let-7(n2853*ts*)/tmC24* | 55% | 0% | n=42 |
| ***let-7(null)/+* genotypes** | | | |
| 25. *lin-41(+); unc-3(e151) let-7(mn112)/tmC24* | 4% | 0% | n=25 |
| 26. *lin-41(tn2128); unc-3(e151) let-7(mn112)/tmC24* | 69% | 3% | n=36 |
| 27. *lin-41(+)/tmC18; unc-3(e151) let-7(mn112)/tmC24* | 4% | 0% | n=25 |
| 28. *lin-41(tn2128)/tmC18; unc-3(e151) let-7(mn112)/tmC24* | 4% | 0% | n=26 |

[a] Animal genotypes are abbreviated for clarity. Balancer chromosome *tmC24* is *let-7(+)* and refers to *tmC24[F23D12.4(tmIs1240) unc-9(tm9719)]*. Balancer chromosome *tmC18* is *lin-41(+)* and refers to *tmC18[dpy-5(tmIs1236)]*.

[b] A strain name is included for *lin-41(+)* genotypes because *let-7(n2853*ts*)* homozygotes from strains DG5827 *let-7/tmC24* and DG6062 *lin-41(+)/tmC18*; *let-7(n2853)/tmC24* exhibited different phenotypes at 15°C. DG6062 likely contains a recessive *let-7* enhancer in the region balanced by *tmC24*; it was created from a strain that had been maintained for several months with *let-7* balanced by *tmC24*.

[c] Egg laying defective or "Egl" animals carried old embryos in their uterus 48 or 72 hours after being selected as L4-stage larvae. Some animals died and were consumed by the larval progeny that hatched inside the uterus. Other animals laid the embryos before they hatched. Both types were scored as "Egl".

[d] "Burst" animals leaked material through the vulva as young adults and died before making many embryos.

[e] Highly variable between replicates (n=3).

**Table S6. *C. elegans* strains used for this study**

| Strain | Genotype |
|---|---|
| N2 | Wild type, Bristol isolate |
| BS553 | *fog-2(oz40)* V |
| CB1370 | *daf-2(e1370)* III |
| CB1372 | *daf-2(e1372)* III |
| DG78 | *smg-1(r861) unc-54(r293)* I |
| DG3501 | *lin-41(ma104)* I |
| DG3712 | *lin-41(tn1492)* I |
| DG3713 | *lin-41(tn1493[C9342351T, acg>ATG, longer uORF])* I |
| DG3714 | *lin-41(tn1494[C9336678T, P942S])* I */ hT2[bli-4(e937) let-?(q782) qIs48]* (I;III) |
| DG3725 | *lin-41(tn1503[C9342482T, acg>ATG, new uORF])* I |
| DG3754 | *lin-41(tn1511)* I |
| DG3755 | *lin-41(tn1512)* I |
| DG3913 | *lin-41(tn1541[gfp::stag::lin-41])* I |
| DG4580 | *lin-41(n2914)* I */ tmC18[dpy-5(tmIs1236 myo-2p::mCherry)]* I |
| DG4727 | *lin-41(tn1892[mScarlet::3xflag::lin-41])* I |
| DG5301 | *lin-41(xe11[3'lcs1&2, c>t]) / tmC18[dpy-5(tmIs1236 myo-2p::mCherry)]* I |
| DG5349 | *lin-41(tn1892[mScarlet::3xflag::lin-41])* I; *fog-2(oz40)* V |
| DG5395 | *lin-41(tn1892[mScarlet::3xflag::lin-41])* I; *lin-29(xe63[gfp::3xflag::lin-29a])* II |
| DG5526 | *lin-41(tn1892[mScarlet::3xflag::lin-41] tn2121[uORFΔ]) / tmC18[dpy-5(tmIs1236 myo-2p::mCherry)]* I |
| DG5528 | *lin-41(tn1892[mScarlet::3xflag::lin-41] tn2121[uORFΔ])* I |
| DG5537 | *tnC1[lin-41(tn1485)]* I |
| DG5542 | *lin-41(tn1892[mScarlet::3xflag::lin-41] tn2121[uORFΔ]) / tmC18[dpy-5(tmIs1236 myo-2p::mCherry)]* I; *lin-29(xe63[gfp::3xflag::lin-29a])* II |
| DG5570 | *lin-41(tn1892[mScarlet::3xflag::lin-41] tn2121[uORFΔ])* I; *fog-2(oz40)* V |
| DG5693 | *lin-41(tn1892[mScarlet::3xflag::lin-41]) / tmC18[dpy-5(tmIs1236 myo-2p::mCherry)]* I*; let-7(n2853*ts*) / tmC24[F23D12.4(tmIs1240 myo-2p::Venus) unc-9(tm9719)]* X |
| DG5694 | *lin-41(tn2128[uORFΔ]) / tmC18[dpy-5(tmIs1236 myo-2p::mCherry)]* I*; let-7(n2853*ts*) / tmC24[F23D12.4(tmIs1240 myo-2p::Venus) unc-9(tm9719)]* X |
| DG5695 | *lin-41(tn1892[mScarlet::3xflag::lin-41])* I*; let-7(n2853*ts*) / tmC24[F23D12.4(tmIs1240 myo-2p::Venus) unc-9(tm9719)]* X |
| DG5696 | *lin-41(tn2128[uORFΔ])* I*; let-7(n2853*ts*) / tmC24[F23D12.4(tmIs1240 myo-2p::Venus) unc-9(tm9719)]* X |
| DG5756 | *lin-41(tn2128[uORFΔ])* I |
| DG5796 | *lin-41(tn1493[acg>ATG, longer uORF])* I; *lin-29(xe63[gfp::3xflag::lin-29a])* II |
| DG5798 | *lin-41(tn1503[acg>ATG, new uORF])* I; *lin-29(xe63[gfp::3xflag::lin-29a])* II |
| DG5804 | *lin-41(ma104)* I; *lin-29(xe63[gfp::3xflag::lin-29a])* II |
| DG5807 | *lin-41(tn2128[uORFΔ])* I; *lin-29(xe63[gfp::3xflag::lin-29a])* II |
| DG5813 | *lin-41(tn1492)* I; *lin-29(xe63[gfp::3xflag::lin-29a]) II* |
| DG5827 | *let-7(n2853*ts*) / tmC24[F23D12.4(tmIs1240 myo-2p::Venus) unc-9(tm9719)]* X |
| DG5835 | *lin-41(tn2128[uORFΔ])* I; *fog-2(oz40)* V |
| DG5855 | *tnEx265[str-1p::gfp]* |
| DG5857 | *lin-41(tn1892[mScarlet::3xflag::lin-41] tn2214[exon0 region2Δ])* I |
| DG5874 | *lin-41(tn2220[exon0Δ]) / tmC18[dpy-5(tmIs1236 myo-2p::mCherry)]* I |

| | |
|---|---|
| DG5877 | *lin-41(tn2220[exon0Δ])* I |
| DG5880 | *lin-41(tn1892[mScarlet::3xflag::lin-41] tn2235[exon0 stop codon])* I |
| DG5891 | *lin-41(tn1892[mScarlet::3xflag::lin-41] tn2232[intron0 splice donor Δ, aaag/g>t])* I |
| DG5892 | *lin-41(tn1892[mScarlet::3xflag::lin-41] tn2233[intron0 splice donor Δ, g/gtgaag>ga])* I |
| DG5922 | *lin-41(tn1892[mScarlet::3xflag::lin-41] tn2241[uORF>lin-29a stem loop1])* I |
| DG5926 | *lin-41(tn1892[mScarlet::3xflag::lin-41] tn2245[5'Δ]) / tmC18[dpy-5(tmIs1236 myo-2p::mCherry)]* I |
| DG5930 | *lin-41(tn1892[mScarlet::3xflag::lin-41] tn2247[uATG>tcc])* I |
| DG5932 | *lin-41(tn1892[mScarlet::3xflag::lin-41] tn2247[uATG>tcc]) / tmC18[dpy-5(tmIs1236 myo-2p::mCherry)]* I |
| DG5944 | *let-7(mn112) unc-3(e151) / tmC24[F23D12.4(tmIs1240 myo-2p::Venus) unc-9(tm9719)]* X |
| DG5945 | *lin-41(tn1892[mScarlet::3xflag::lin-41] tn2250[uORF>mutant lin-29a stem loop1])* I |
| DG5963 | *lin-41(tn1892[mScarlet::3xflag::lin-41] tn2252[exon0 $1^{st}$ acg>tcc])* I |
| DG6003 | *lin-41(tn1892[mScarlet::3xflag::lin-41] tn2285[exon0 region1Δ])* I |
| DG6033 | *lin-41(tn2128[uORFΔ] tn2293[exon0 region3Δ])* I |
| DG6040 | *lin-41(tn1892[mScarlet::3xflag::lin-41]) unc-54(r293)* I |
| DG6042 | *smg-2(r908) lin-41(tn1892[mScarlet::3xflag::lin-41]) unc-54(r293)* I |
| DG6043 | *lin-41(tn1892[mScarlet::3xflag::lin-41] tn2295[exon0 region3Δ])* I |
| DG6046 | *lin-41(tn2220[exon0Δ]) / tmC18[dpy-5(tmIs1236 myo-2p::mCherry]* I; *lin-29(xe63[gfp::3xflag::lin-29a]) II* |
| DG6049 | *lin-41(tn2297[exon0 region4Δ (includes uORF)])* I |
| DG6061 | *lin-41(tn2220[exon0Δ]) / tmC18[dpy-5(tmIs1236 myo-2p::mCherry)]* I; *let-7(n2853ts) / tmC24[F23D12.4(tmIs1240 myo-2p::Venus) unc-9(tm9719)]* X |
| DG6062 | *lin-41(+) / tmC18[dpy-5(tmIs1236 myo-2p::mCherry]* I; *let-7(n2853ts) / tmC24[F23D12.4(tmIs1240 myo-2p::Venus) unc-9(tm9719)]* X |
| DG6094 | *lin-41(tn2128[uORFΔ]) / tmC18[dpy-5(tmIs1236 myo-2p::mCherry)]* I*; let-7(mn112) unc-3(e151) / tmC24[F23D12.4(tmIs1240 myo-2p::Venus) unc-9(tm9719)]* X |
| DG6095 | *lin-41(+) / tmC18[dpy-5(tmIs1236 myo-2p::mCherry)]* I*; let-7(mn112) unc-3(e151) / tmC24[F23D12.4(tmIs1240 myo-2p::Venus) unc-9(tm9719)]* X |
| DG6107 | *lin-41(tn1892[mScarlet::3xflag::lin-41] tn2121[uORFΔ]) / tmC18[dpy-5(tmIs1236 myo-2p::mCherry)]* I; *let-7(n2853ts) / tmC24[F23D12.4(tmIs1240) unc-9(tm9719)]* X |
| DG6108 | *lin-41(+) / tmC18[dpy-5(tmIs1236 myo-2p::mCherry]* I; *let-7(n2853ts) / tmC24[F23D12.4(tmIs1240 myo-2p::Venus) unc-9(tm9719)]* X |
| DG6112 | *lin-41(tn1892[mScarlet::3xflag::lin-41] tn2330[intron0Δ]) / tmC18[dpy-5(tmIs1236 myo-2p::mCherry)]* I |
| DG6119 | *lin-41(tn2333[intron0Δ]) / tmC18[dpy-5(tmIs1236 myo-2p::mCherry)]* I |
| DG6151 | *lin-41(tn1892[mScarlet::3xflag::lin-41] tn2330[intron0Δ] tn2351[uATG > tcc]) / tmC18[dpy-5(tmIs1236 myo-2p::mCherry)]* I |
| DG6153 | *lin-41(tn1892 [mScarlet::3xflag::lin-41] tn2330[intron0Δ] tn2351[uATG>tcc])* |
| DG6155 | *lin-41tn2333[intron0Δ] tn2353[uATG>tcc]) / tmC18[dpy-5(tmIs1236 myo-2p::mCherry)]* I |

| | |
|---|---|
| DG6182 | *lin-41(tn1892[mScarlet::3xflag::lin-41])* I; *mgIs49(mlt-10p::gfp)* IV |
| DG6183 | *lin-41(tn1892[mScarlet::3xflag::lin-41] tn2121[uORFΔ]* I; *mgIs49(mlt-10p::gfp)* IV |
| DG6184 | *lin-41(tn1892[mScarlet::3xflag::lin-41] tn2247[uATG>tcc])* I; *mgIs49(mlt-10p::gfp)* IV |
| DG6185 | *lin-41(tn2128[uORFΔ])* I; *mgIs49(mlt-10p::gfp)* IV |
| DG6186 | *lin-41(tn2220[exon0Δ])* I; *mgIs49(mlt-10p::gfp) IV* |
| DG6187 | *lin-41(tn1892[mScarlet::3xflag::lin-41] tn2330[intron0Δ]) / tmC18[dpy-5(tmIs1236 myo-2p::mCherry)]* I; *mgIs49(mlt-10p::gfp)* IV |
| DG6188 | *lin-41(tn1892[mScarlet::3xflag::lin-41] tn2330[intron0Δ] tn2351[uATG>tcc]) / tmC18[tmIs1236(myo-2p::mCherry)]* I; *mgIs49(mlt-10p::gfp)* IV |
| DG6219 | *lin-41(tn2220[exon0Δ]) / tmC18[dpy-5(tmIs1236 myo-2p::mCherry)]* I *; daf-7(e1372)* III |
| DG6247 | *lin-41(tn2220[exon0Δ]) / tmC18[dpy-5(tmIs1236 myo-2p::mCherry)]* I*; daf-7(e1372)* III; *syIs601[ets-10p::GFP + ofm-1p::RFP]* |
| DG6249 | *daf-7(e1372)* III; *syIs601[ets-10p::GFP + ofm-1p::RFP]* |
| DG6250 | *daf-2(e1370)* III; *syIs601[ets-10p::GFP + ofm-1p::RFP]* |
| DG6261 | *lin-41(tn2220[exon0Δ]) / tmC18[dpy-5(tmIs1236 myo-2p::mCherry)]* I*; syIs601[ets-10p::GFP + ofm-1p::RFP]* |
| DG6304 | *lin-41(tn1892[mScarlet::3xflag::lin-41] tn2121[uORF*D*])* I; *let-7(n2853*ts*) / tmC24[F23D12.4(tmIs1240 myo-2p::Venus) unc-9(tm9719)]* X |
| DG6305 | *let-7(n2853*ts*) / tmC24[F23D12.4(tmIs1240 myo-2p::Venus) unc-9(tm9719)]* X (derived from DG6062) |
| DG6306 | *lin-41(tn2220[exon0Δ])* I; *let-7(n2853*ts*) / tmC24[F23D12.4(tmIs1240 myo-2p::Venus) unc-9(tm9719)]* X |
| DG6311 | *lin-41(tn2220[exon0Δ]) / tmC18[dpy-5(tmIs1236 myo-2p::mCherry)]* I*; daf-2(e1370)* III; *syIs601[ets-10p::GFP + ofm-1p::RFP]* |
| DG6335 | *lin-41(tn2454[exon0 region1,2Δ])* I |
| DG6337 | *smg-2(r908) lin-41(tn1503) unc-54(r293)* I |
| DG6338 | *smg-2(r908) lin-41(tn1493) unc-54(r293)* I |
| DG6355 | *lin-41(tn2454[exon0 region1,2Δ] tn2463[uATG>tcc])* I |
| DG6360 | *lin-41(tn2454[exon0 region1,2Δ] tn2466[11bp uATG Δ])* / *tmC18[dpy-5(tmIs1236 myo-2p::mCherry)]* I |
| DG6361 | *lin-41(tn2454[exon0 region1,2Δ] tn2466[11bp uATG Δ])* I |
| DG6362 | *lin-41(tn2454[exon0 region1,2Δ] tn2467[23bp uATG Δ])* / *tmC18[dpy-5(tmIs1236 myo-2p::mCherry)]* I |
| DG6363 | *lin-41(tn2454[exon0 region1,2Δ] tn2467[23bp uATG Δ])* I |
| FX30167 | *tmC18[dpy-5(tmIs1200 myo-2p::Venus)]* I |
| FX30168 | *tmC18[dpy-5(tmIs1236 myo-2p::mCherry)]* I |
| FX30194 | *tmC24[F23D12.4(tmIs1240 myo-2p::Venus) unc-9(tm9719)]* X |
| FX30240 | *tmC24[F23D12.4(tmIs1240 myo-2p::Venus)]* X |
| FX30252 | *tmC24[F23D12.4(tmIs1240 myo-2p::Venus) unc-9(tm9719)]* X; *tmEx4950[unc-9(+) + vha-6p::gfp]* |
| GR1395 | *mgIs49(mlt-10p::gfp)* IV |
| HW1826 | *lin-29(xe63[gfp::3xflag::lin-29a])* II |
| JK4862 | *glp-1(q46)* III / *hT2[bli-4(e937) let-?(q782) qIs48(myo-2p::gfp)]* (I;III) |
| PS8457 | *syIs601[ets-10p::GFP + ofm-1p::RFP]* |

| | |
|---|---|
| TR1421 | *smg-2(r908) unc-54(r293)* I |

**Table S7** (Related to Fig. 9). **Sequence of smFISH probes and transcript count data.**

**Table S8 (**Related to Figs 2, 4, 7- 9, S6, S7, S9, Tables S4 and S5**). *lin-41* phenotype data and statistical analyses.**

### LEGENDS TO SUPPLEMENTAL FIGURES

**Fig. S1** (Related to Fig. 2). **RT-PCR analysis of *lin-41* transcripts.** An ethidium bromide-stained 2% agarose gel showing the detection of *lin-41* long and *lin-41* short transcripts in the wild type and mutants. Approximately 200 ng of purified RT-PCR products that were used for exome sequencing were analyzed. The results of exome sequencing of these samples are shown in Table S1. As noted in Materials and Methods, because somatic *lin-41* transcripts are apparently of low abundance in adults, 10 50 µl PCR reactions were conducted and pooled to get sufficient quantities of cDNA for exome sequencing from *glp-1(q46)* adults. The PCR products shown are representative of at least three replicates.

**Fig. S2.** (Related to Fig. 2). **Analysis of ribosomal footprinting data.** (A,B) Ribosomal footprinting data from Hendriks et al. (2014) and Aeschimann et al. (2017) were reanalyzed to assess ribosome-protected fragments in *lin-41* exon 0. Data are from the wild type and *let-7(n2853*ts*)* at the indicated times post-hatch at 25°C are shown. In order to compare samples, the number of ribosome-protected fragments in an experiment was normalized according to the total number of ribosome-protected fragments mapping to *lin-41* exons in that experiment. (A) Normalized reads across *lin-41*. Arrowheads indicate a small peak of ribosome-protected fragments within exon 0. (B) A zoom-in comparison of exon 0 and exon 1. Exon 0 is shown and the *lin-41* uORF is indicated by a black triangle. See Table S2 for the tabulation of data.

**Fig. S3** (Related to Fig. 4). ***lin-41* exon 0 gain-of-function mutants inappropriately activate a molting program late in the adult stage.** (A-H) Overlay of DIC and GFP fluorescence images of the indicated strains showing the activation of *mlt-10p::gfp* at 72-73 hr after feeding L1 larvae, which were isolated by alkaline hypochlorite treatment and allowed to hatch in the absence of food. The number of GFP-positive worms out of the number scored is shown in the lower left corner of each panel. Images outlined in magenta show animals with the *lin-41(tn1892[mScarlet::lin-41])* genetic background. Bars, 200 µm.

**Fig. S4** (Related to Fig. 4). **Deletion of the *lin-41* uORF causes a gain-of-function phenotype in the *lin-41(tn1892[mScarlet::lin-41])* genetic background.** (A-D) DIC micrographs of the indicated genotypes showing that *lin-41(tn1892[mScarlet::lin-41] tn2121[uORFΔ])* males exhibit a Lep phenotype (D). Bars, 20 μm.

**Fig. S5** (Related to Fig. 4). **RT-PCR analysis of *lin-41* transcripts in the *lin-41(tn1892[mScarlet::lin-41])* genetic background.** An ethidium bromide-stained 2% agarose gel showing the detection of *lin-41* long and *lin-41* short transcripts. Approximately 200 ng of purified RT-PCR products were analyzed. The results of exome sequencing of these samples are shown in Table S1. *lin-41(tn1892[mScarlet::lin-41] tn2245 [5'Δ])* produces a novel *lin-41* long transcript (indicated by an asterisk) owing to the deletion of the SL1 splice acceptor sequence and the use of a cryptic SL1 splice acceptor site. The indicated PCR products are non-specific (ns) as they do not originate from the *lin-41* locus. These non-specific products were most abundant in the *lin-41(tn1892[mScarlet::lin-41] tn2245 [5'Δ])* sample potentially owing to a low abundance of *lin-41* transcripts in this mutant. This observation accounts for the fewer number of reads for this mutant in the exome sequencing results (Table S1). As noted in Materials and Methods, 20 50 μl PCR reactions were needed to get sufficient quantities of cDNA from *lin-41(tn1892 tn2245)* L1-L3 larvae for exome sequencing. The PCR products shown are representative of at least three replicates.

**Fig. S6** (Related to Fig. 7). **Quantification of mScarlet::LIN-41 levels.** (A) The *smg-2(r908)* mutation does not increase expression of mScarlet::LIN-41. Background-corrected mScarlet::LIN-41 fluorescence levels in DG4727 *lin-41(tn1892[mScarlet::lin-41])* and DG6042 *smg-2(r908) lin-41(tn1892[mScarlet::lin-41]) unc-54(r293)* L4 larvae, measured in the head region. The *unc-54(r293)* mutation is suppressed by *smg-2(r908)*. Dots are data points, center horizontal lines indicate the median, boxes indicate the interquartile range and whiskers encompass 80% of the data. *P*=0.278, *t*-test with

Welch's correction. (B) The *lin-41* uORF and Let-7 coordinately regulate mScarlet::LIN-41 expression. Background-corrected mScarlet::LIN-41 fluorescence levels measured in the head region of L4 larvae at 20°C of animals with the following genotypes from the indicated strains: DG4727 *lin-41(tn1892[mScarlet::lin-41])*, DG5930 *lin-41(tn1892[mScarlet::lin-41] tn2247[uATG>tcc])*, *lin-41(tn1892[mScarlet::lin-41] tn2121[uORFΔ])* from DG5526, *lin-41(tn1892[mScarlet::lin-41]; let-7(n2853*ts*)* from DG5695, and *lin-41(tn1892[mScarlet::lin-41] tn2121[uORFΔ]); let-7(n2853*ts*)* from DG6304. Dots are data points, center horizontal lines indicate the median, boxes indicate the interquartile range and whiskers encompass 80% of the data. Brackets indicate a statistical comparison between the indicated genotypes. Significance values that are directly above a distribution indicate a comparison to animals with the *lin-41(tn1892)* control genotype. Statistical analyses utilized an ANOVA with a post-hoc Games-Howell test. Raw data and statistics are in Table S8.

**Fig. S7** (Related to Fig. 7). **Increased mScarlet::LIN-41 levels in *lin-41(tn1892[mScarlet::lin-41[5'Δ])* mutants.** (A,B) Fluorescence micrographs mScarlet::LIN-41 expression (left panels) L4 larvae of the indicated genotypes and shown overlaid with the DIC images (right panels). The boxed insets at the right are closeups (two-fold magnified) showing the vulval region (V), which was used for staging the animals. The relative fluorescence levels are indicated in the left panels. Bars, 50 μm. (C) Background-corrected mScarlet::LIN-41 fluorescence levels measured in the head region of L4 larvae at 20°C of animals with the following genotypes from the indicated strains: DG4727 *lin-41(tn1892[mScarlet::lin-41])* and *lin-41(tn1892[mScarlet::lin-41] tn2245[5'Δ])* from DG5926. Dots are data points, center horizontal lines indicate the median, boxes indicate the interquartile range and whiskers encompass 80% of the data. Statistical analysis employed a *t*-test with Welch's correction, $P$=1.03x$10^{-11}$. Raw data and statistics are in Table S8.

**Fig. S8** (Related to Fig. 9**). Intron 0 promotes *lin-41* transcript accumulation in developing oocytes.** (A-C) *lin-41* smFISH (yellow on the left, grayscale on the right) in dissected adult hermaphrodite gonads from the wild type (A), *lin-41(tn2333[intron0Δ])* (B) and *lin-41(tn2333[intron0Δ] tn2353[uATG>tcc])* mutants (C). WGA staining (magenta) was used to delineate oocyte boundaries. DNA is cyan. (D) *lin-41* smFISH quantification in diakinesis-stage oocytes in the wild type and *lin-41(tn2333[intron0Δ] tn2353[uATG>tcc])* mutants. *lin-41(tn2333[intron0Δ])* oocytes were too disorganized to permit reliable smFISH quantification on a per-oocyte basis. Dots are data points, boxplots with whiskers show the median, interquartile range and 80% of the data points. Statistical analysis employed a *t*-test with Welch's correction, ****P<0.0001. Raw data and statistics are in Table S7.

**Fig. S9** (Related to Fig. 9). **Exon 0 and its embedded uORF affect mScarlet::LIN-41 expression in the germline.** mScarlet::LIN-41 expression in *lin-41[tn1892[mScarlet::lin-41])* (A and C-D), *lin-41[tn1892[mScarlet::lin-41][tn2330[intron0Δ])* (B-D and E) and *lin-41[tn1892[mScarlet::lin-41] [tn2330[intron0Δ] tn2351[uATG>tcc])* (F) adult hermaphrodites. mScarlet::LIN-41 images in (A,B) are overlaid with the DIC channel and were collected using different settings from the images in (E,F). Graphs quantify the spatio-temporal (C) and mean mScarlet::LIN-41 expression (D) during oogenesis in animals imaged as in (A,B). Dots are individual data points showing per gonad measurements, center horizontal lines indicate the median, boxes indicate the interquartile range and whiskers encompass all the data in (D). Statistical analysis employed a *t*-test with Welch's correction. Arrowheads indicate small abnormal oocytes in *lin-41[tn1892[mScarlet::lin-41][tn2330[intron0Δ])* mutants (B,E). Day 1 adults, 24 h post L4 were analyzed in (A-D), and young day 1 adults, 16 h post L4 were analyzed in (E,F). Bars, 50 µm. Raw data and statistics are in Table S8.

**Fig. S10** (Related to Fig. 10). **Exon 0 is conserved in Caenorhabditid nematodes.** Sequence alignment of exon 0 in the indicated nematode species. Inverted triangles indicate the beginning (filled triangle) and end (open triangle) of exon 0 in *Caenorhabditis elegans*. SL1 is added after the sequence 5'-TTTCAG-3' (open triangle) and the 5'-GT-3' splice donor of intron 0 is indicated by the filled triangle. Regions 1–4 of exon 0 are indicated as is the *lin-41* uORF coding sequence.

**Fig. S11** (Related to Discussion). **Computational model for the coordinate regulation of LIN-41 by Let-7 and exon 0.** (A) A three-dimensional plot of LIN-41 protein levels as a function of *let-7* activity and exon 0 strength using a rheostat-based model (Equation 1) at the L4 larval stage when *lin-41* activity is downregulated in the wild type. Movie 1 shows a rotational animation of this plot. (B) A planar projection of the plot shown in (A). A $P_{LIN-41}$ activity level of 1 is assigned to the L4 larval stage in the wild type. Estimated activity thresholds at the L4 larval stage for WT (~1–1.8), egg-laying defective (~1.8–2.5), and lethal bursting phenotypes (~2.5–4.8) are indicated by contours corresponding to $P_{LIN-41}$=01.8 (solid line) and $P_{LIN-41}$=2.5 (dashed line).

**Movie 1** (Related to Fig. S11). **Model for the coordinate regulation of LIN-41 by Let-7 and exon 0.** Rotational animation of the three-dimensional plot showing LIN-41 protein levels as a function of *let-7* activity and exon 0 strength at the L4 larval stage when *lin-41* activity is downregulated in the wild type.

**Fig. S1**

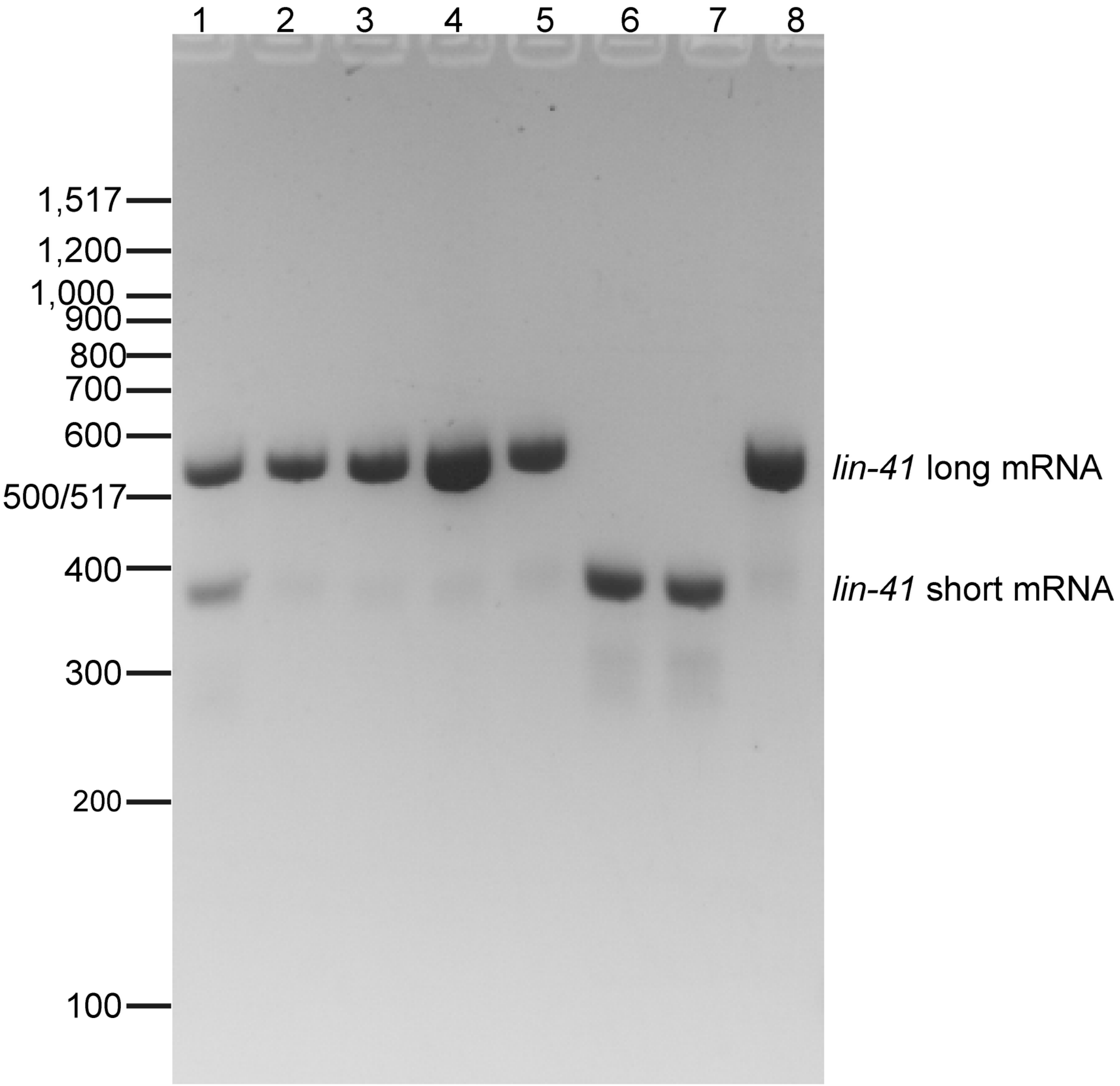


1. WT adult
2. WT L1
3. WT L3
4. *lin-41(tn2333 [intron 0Δ])* adult
5. *lin-41 (tn2333 [intron 0Δ] tn2353 [uATC>tcc])* adult
6. *lin-41(tn2220 [exon 0Δ])* adult
7. *lin-41(tn2220 [exon 0Δ])*L3
8. *glp-1(q46)* adult

Fig. S2

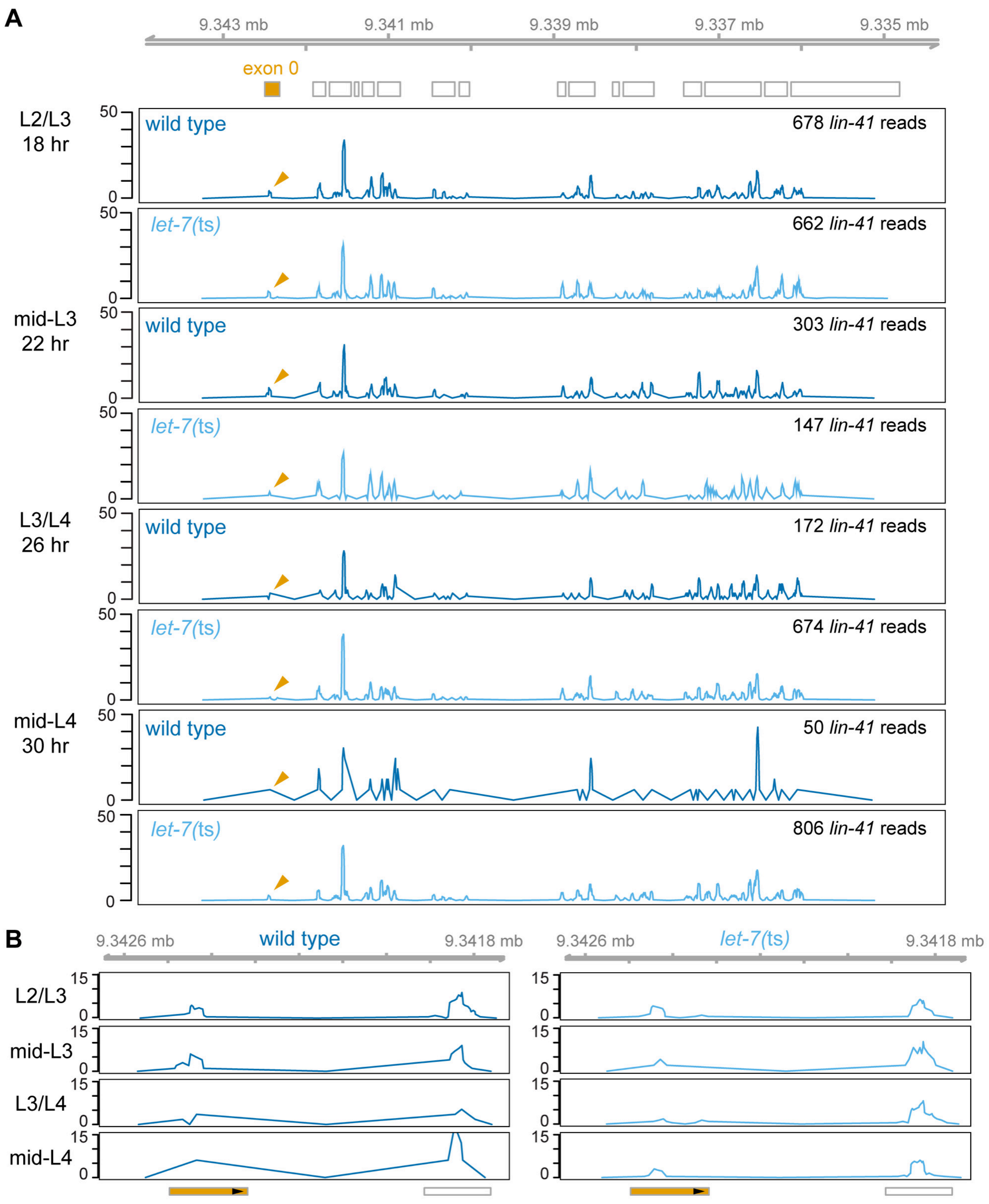
A
9.343 mb
9.341 mb
9.339 mb
9.337 mb
9.335 mb
exon 0
L2/L3
18 hr
wild type
678 lin-41 reads
let-7(ts)
662 lin-41 reads
mid-L3
22 hr
wild type
303 lin-41 reads
let-7(ts)
147 lin-41 reads
L3/L4
26 hr
wild type
172 lin-41 reads
let-7(ts)
674 lin-41 reads
mid-L4
30 hr
wild type
50 lin-41 reads
let-7(ts)
806 lin-41 reads
50
0
B
9.3426 mb
wild type
9.3418 mb
9.3426 mb
let-7(ts)
9.3418 mb
L2/L3
mid-L3
L3/L4
mid-L4
15
0

### Fig. S3

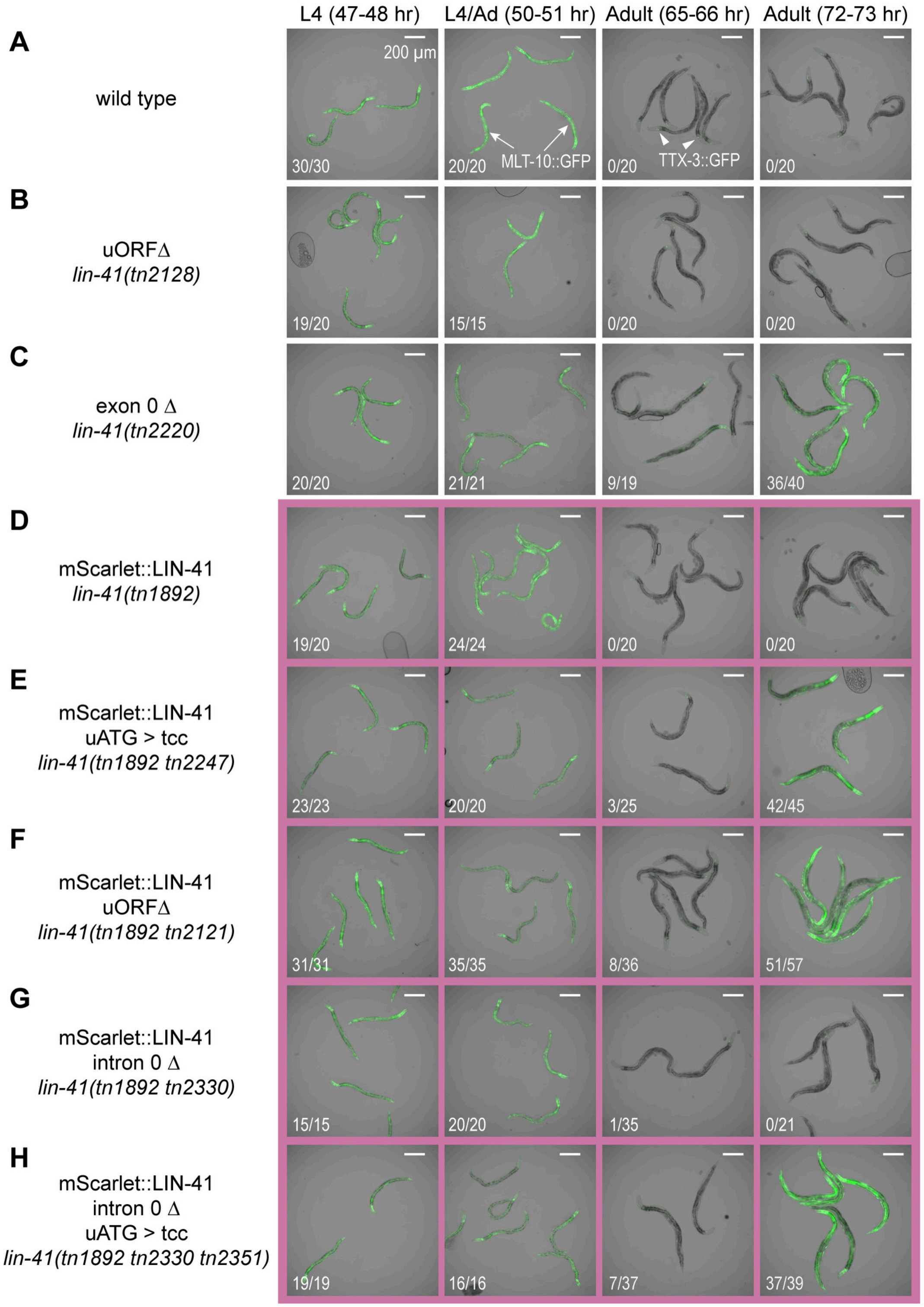
L4 (47-48 hr)
L4/Ad (50-51 hr)
Adult (65-66 hr)
Adult (72-73 hr)
A
wild type
200 µm
30/30
20/20
MLT-10::GFP
0/20
TTX-3::GFP
0/20
B
uORFΔ
lin-41(tn2128)
19/20
15/15
0/20
0/20
C
exon 0 Δ
lin-41(tn2220)
20/20
21/21
9/19
36/40
D
mScarlet::LIN-41
lin-41(tn1892)
19/20
24/24
0/20
0/20
E
mScarlet::LIN-41
uATG > tcc
lin-41(tn1892 tn2247)
23/23
20/20
3/25
42/45
F
mScarlet::LIN-41
uORFΔ
lin-41(tn1892 tn2121)
31/31
35/35
8/36
51/57
G
mScarlet::LIN-41
intron 0 Δ
lin-41(tn1892 tn2330)
15/15
20/20
1/35
0/21
H
mScarlet::LIN-41
intron 0 Δ
uATG > tcc
lin-41(tn1892 tn2330 tn2351)
19/19
16/16
7/37
37/39

**Fig. S4**

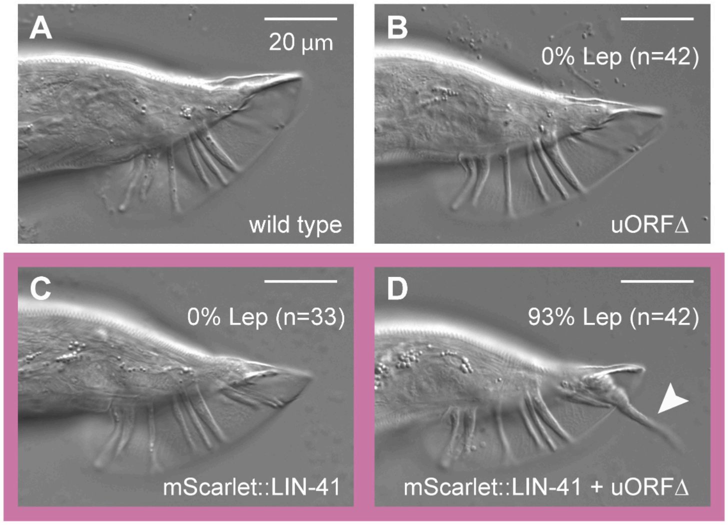

A
20 µm
wild type
B
0% Lep (n=42)
uORFΔ
C
0% Lep (n=33)
mScarlet::LIN-41
D
93% Lep (n=42)
mScarlet::LIN-41 + uORFΔ

**Fig. S5**

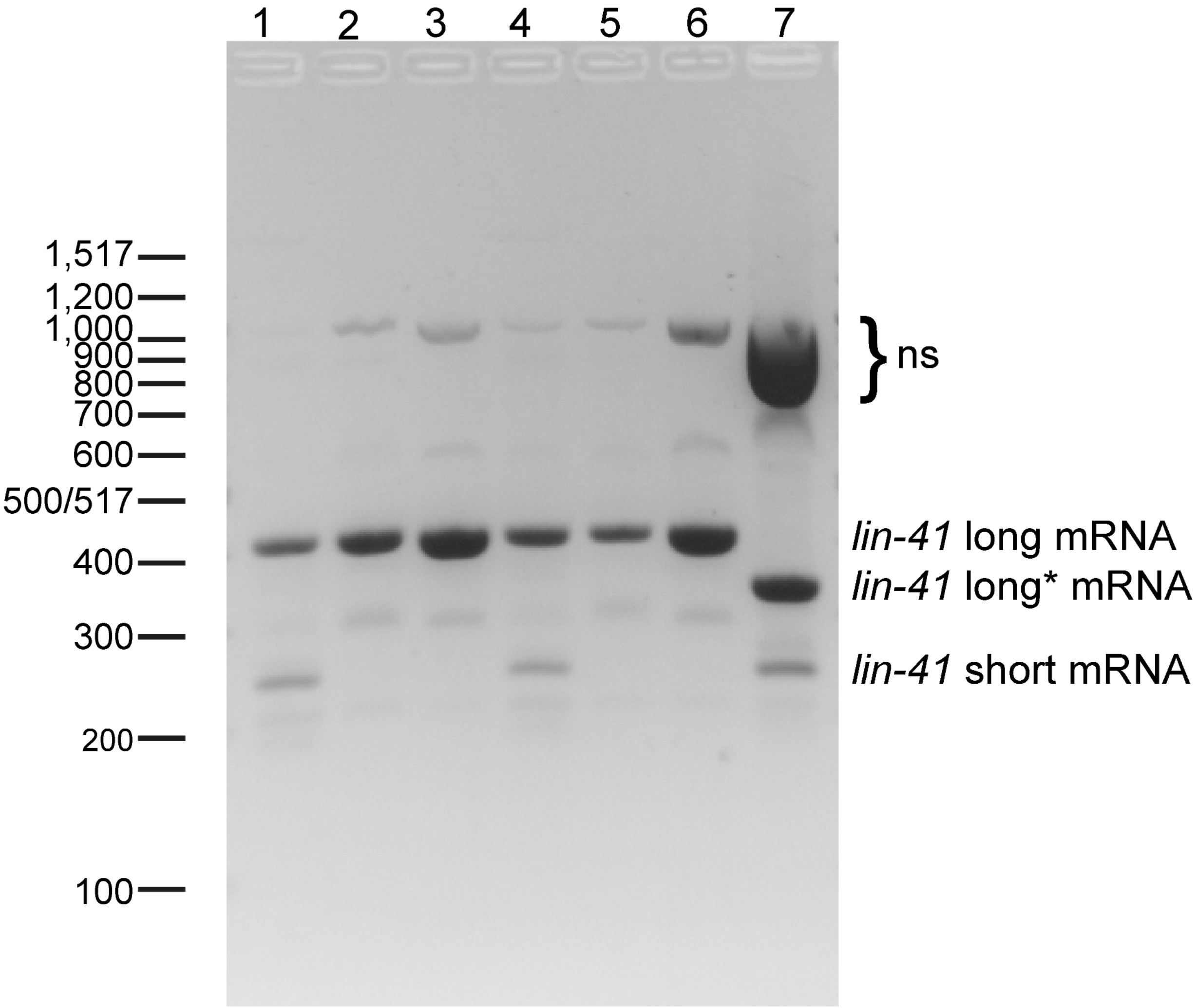


1. *lin-41(tn1892[mScarlet::lin-41])* adult
2. *lin-41(tn1892[mScarlet::lin-41])* L1
3. *lin-41(tn1892[mScarlet::lin-41])* L3
4. *lin-41(tn1892[mScarlet::lin-41] tn2247[uATG to tcc])* adult
5. *lin-41(tn1892[mScarlet::lin-41] tn2247[uATG to tcc])* L1
6. *lin-41(tn1892[mScarlet::lin-41] tn2247[uATG to tcc])* L3
7. *lin-41(tn1892[mScarlet::lin-41] tn2245[5'Δ])* L1-L3

**Fig. S6**

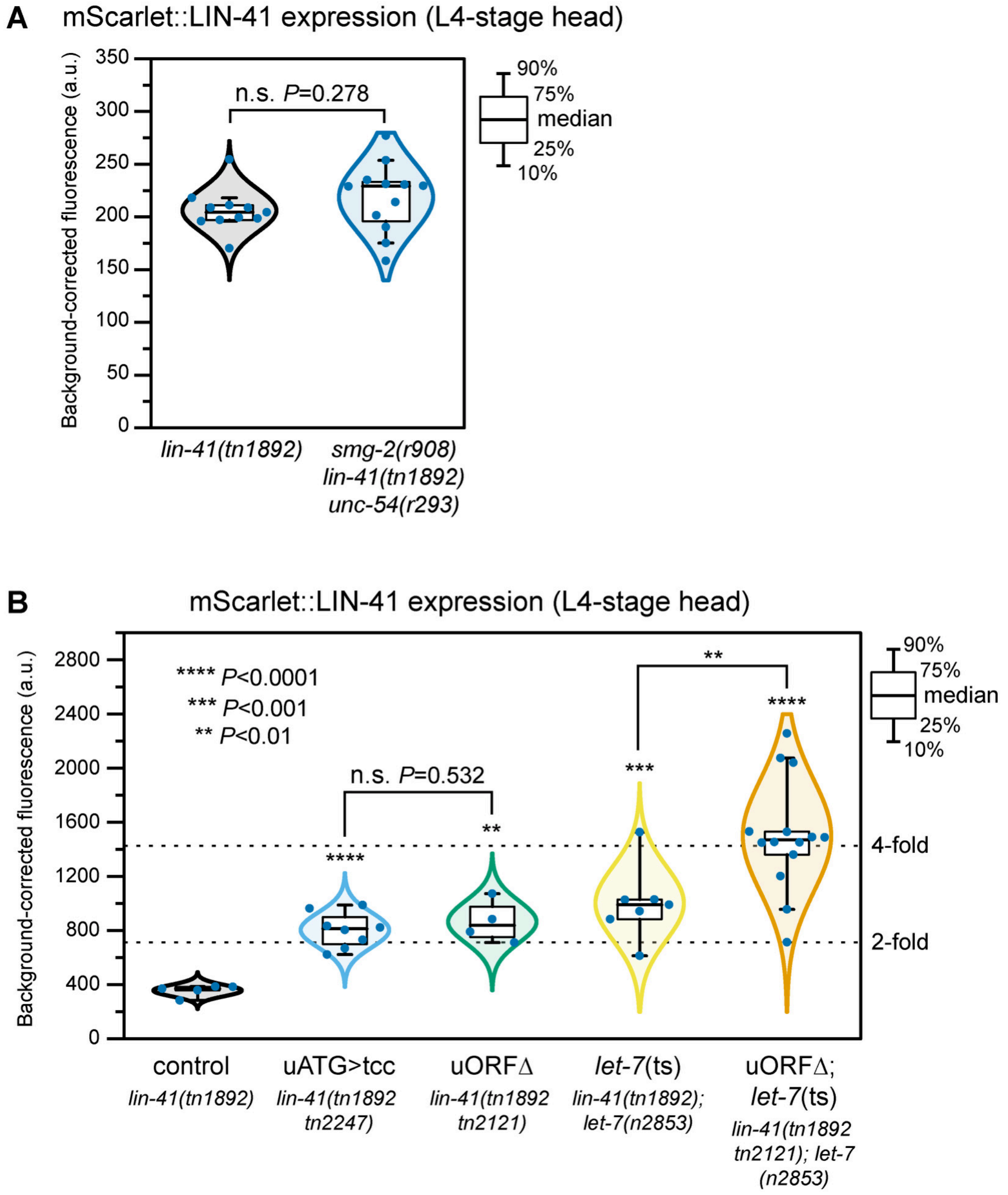
A
mScarlet::LIN-41 expression (L4-stage head)
Background-corrected fluorescence (a.u.)
350
300
250
200
150
100
50
0
n.s. P=0.278
90%
75%
median
25%
10%
lin-41(tn1892)
smg-2(r908)
lin-41(tn1892)
unc-54(r293)
B
mScarlet::LIN-41 expression (L4-stage head)
Background-corrected fluorescence (a.u.)
2800
2400
2000
1600
1200
800
400
0
**** P<0.0001
*** P<0.001
** P<0.01
n.s. P=0.532
****
**
***
**
****
4-fold
2-fold
90%
75%
median
25%
10%
control
lin-41(tn1892)
uATG>tcc
lin-41(tn1892 tn2247)
uORFΔ
lin-41(tn1892 tn2121)
let-7(ts)
lin-41(tn1892); let-7(n2853)
uORFΔ; let-7(ts)
lin-41(tn1892 tn2121); let-7 (n2853)

Fig. S7

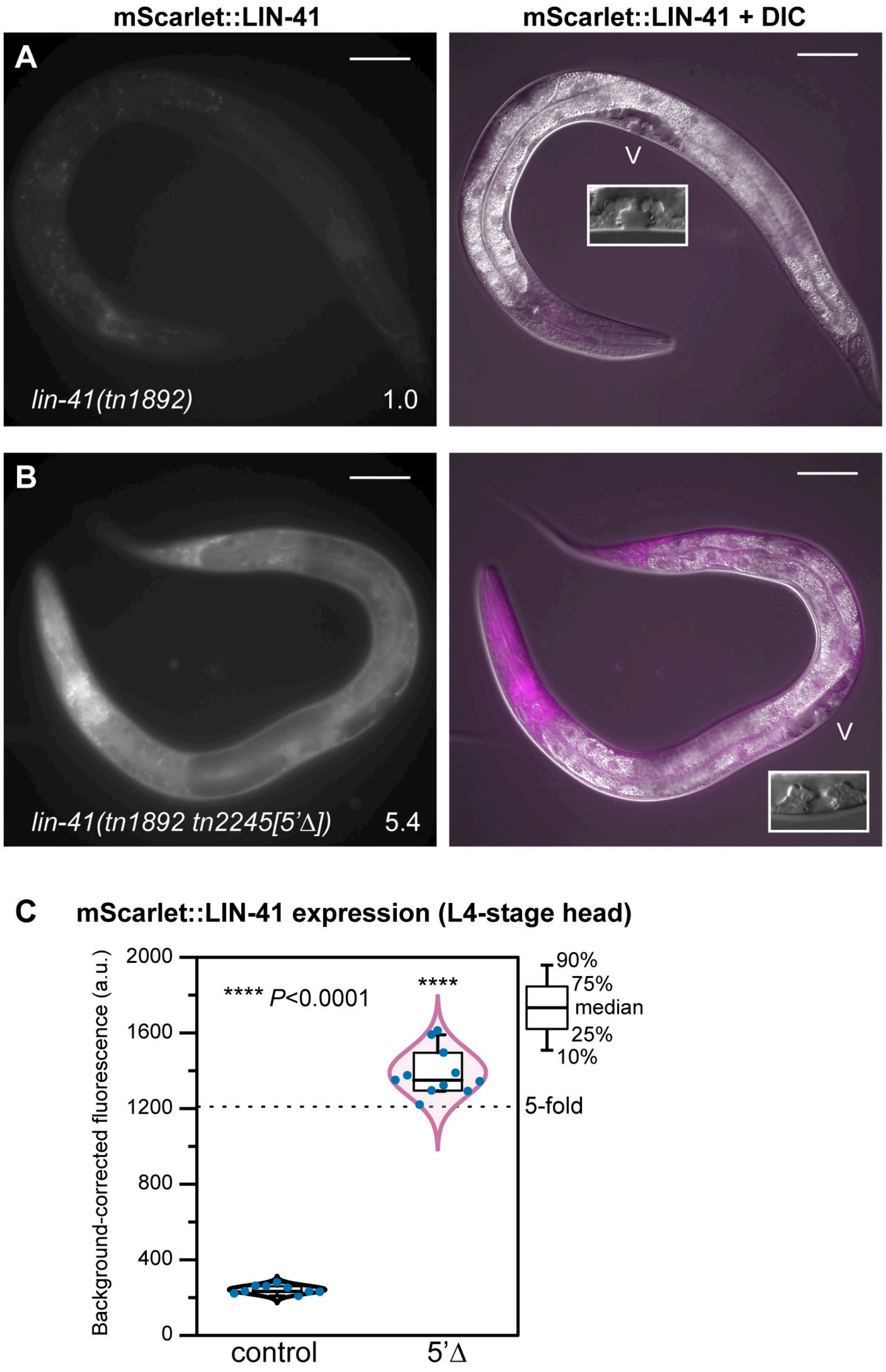

mScarlet::LIN-41
mScarlet::LIN-41 + DIC
A
lin-41(tn1892)
1.0
V
B
lin-41(tn1892 tn2245[5'Δ])
5.4
V
C
mScarlet::LIN-41 expression (L4-stage head)
Background-corrected fluorescence (a.u.)
2000
1600
1200
800
400
0
**** P<0.0001
****
5-fold
90%
75%
median
25%
10%
control
5'Δ

**Fig. S8**

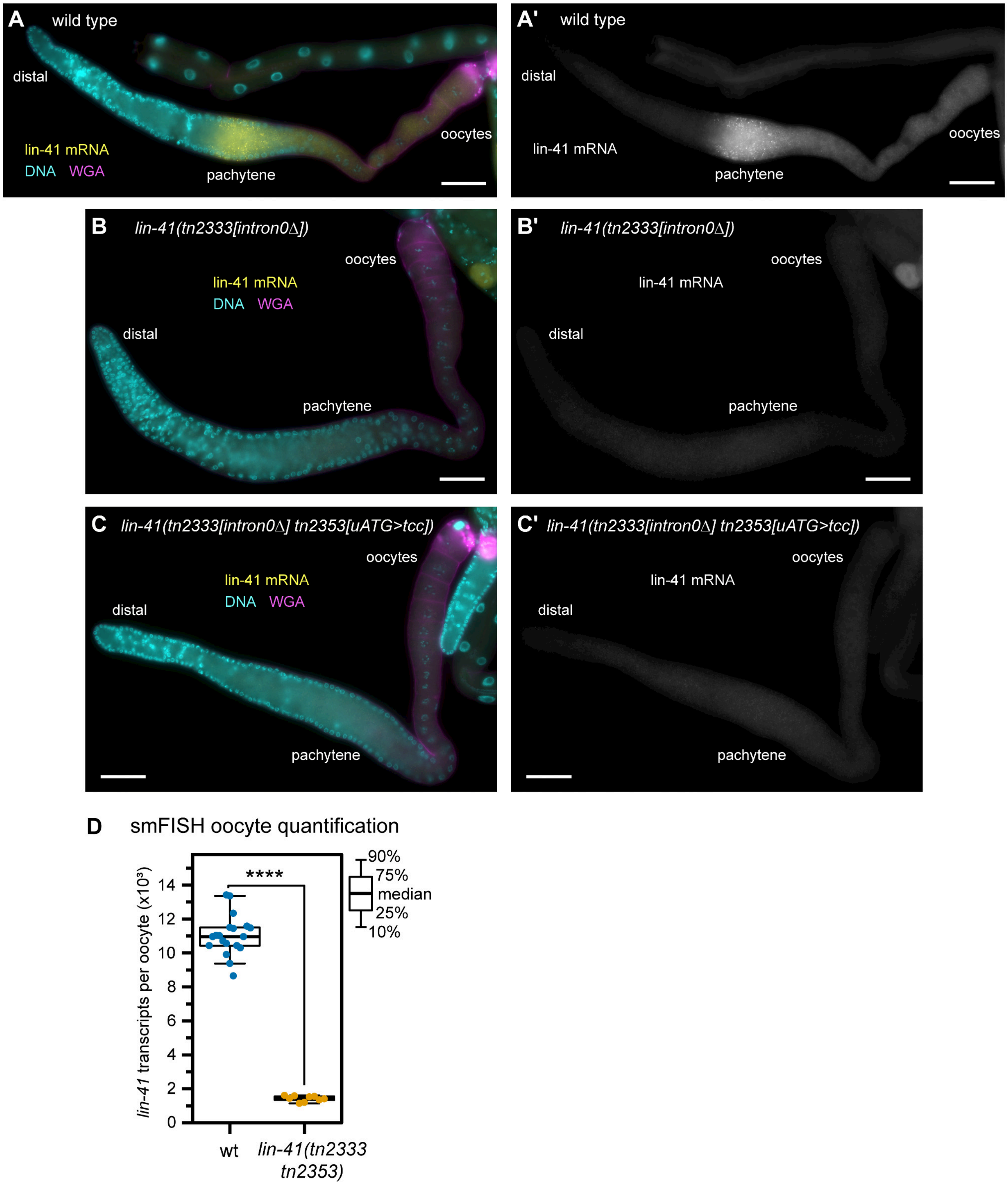
A
wild type
distal
lin-41 mRNA
DNA
WGA
pachytene
oocytes
A'
wild type
distal
lin-41 mRNA
pachytene
oocytes
B
lin-41(tn2333[intron0Δ])
oocytes
lin-41 mRNA
DNA
WGA
distal
pachytene
B'
lin-41(tn2333[intron0Δ])
oocytes
lin-41 mRNA
distal
pachytene
C
lin-41(tn2333[intron0Δ] tn2353[uATG>tcc])
oocytes
lin-41 mRNA
DNA
WGA
distal
pachytene
C'
lin-41(tn2333[intron0Δ] tn2353[uATG>tcc])
oocytes
lin-41 mRNA
distal
pachytene
D
smFISH oocyte quantification
lin-41 transcripts per oocyte (x10³)
0
2
4
6
8
10
12
14
****
90%
75%
median
25%
10%
wt
lin-41(tn2333 tn2353)

**Fig. S9**

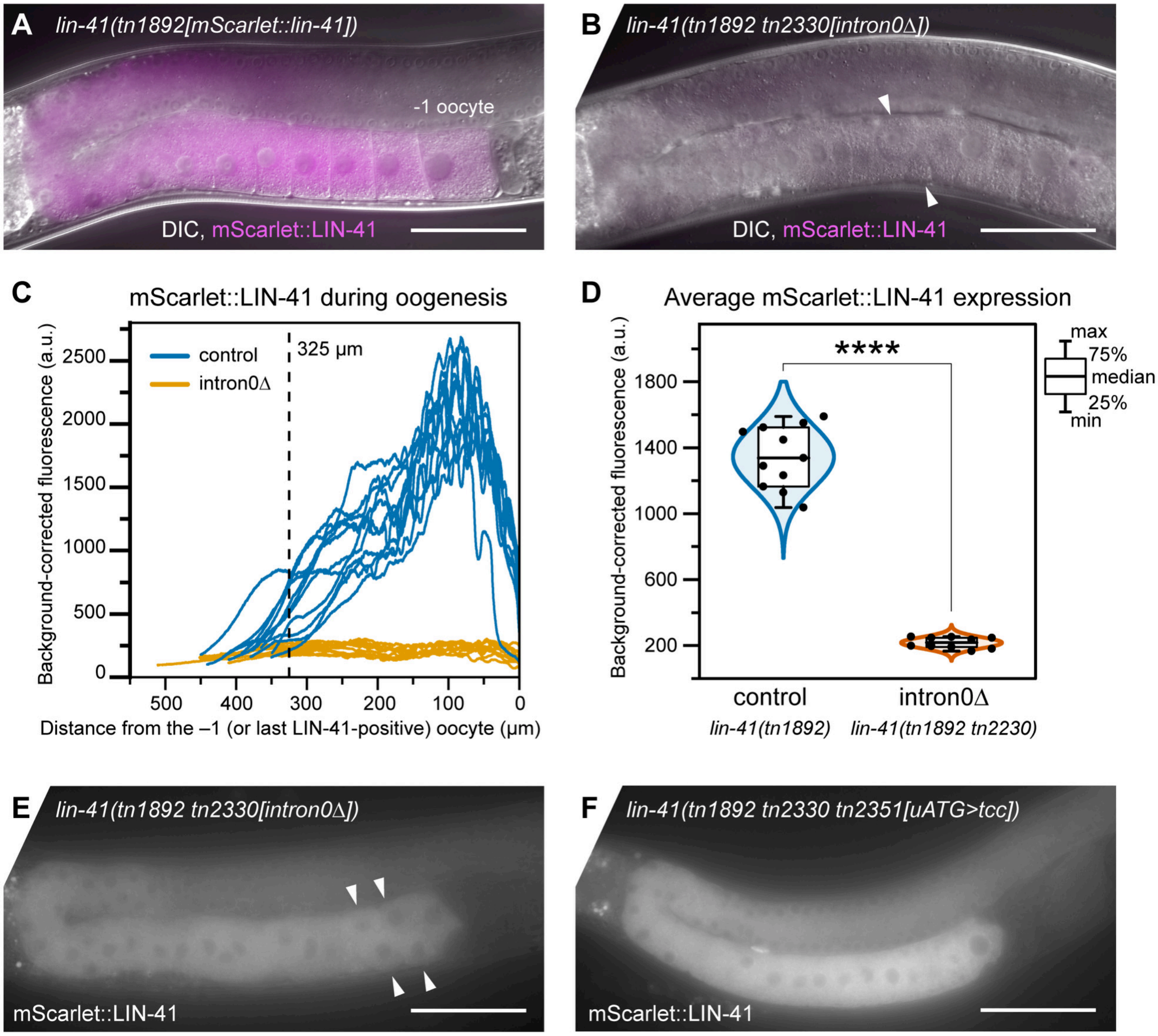

A
lin-41(tn1892[mScarlet::lin-41])
-1 oocyte
DIC, mScarlet::LIN-41
B
lin-41(tn1892 tn2330[intron0Δ])
DIC, mScarlet::LIN-41
C
mScarlet::LIN-41 during oogenesis
Background-corrected fluorescence (a.u.)
control
intron0Δ
325 μm
2500
2000
1500
1000
500
0
500 400 300 200 100 0
Distance from the –1 (or last LIN-41-positive) oocyte (μm)
D
Average mScarlet::LIN-41 expression
Background-corrected fluorescence (a.u.)
****
1800
1400
1000
600
200
max
75%
median
25%
min
control
lin-41(tn1892)
intron0Δ
lin-41(tn1892 tn2230)
E
lin-41(tn1892 tn2330[intron0Δ])
mScarlet::LIN-41
F
lin-41(tn1892 tn2330 tn2351[uATG>tcc])
mScarlet::LIN-41

**Fig. S10**

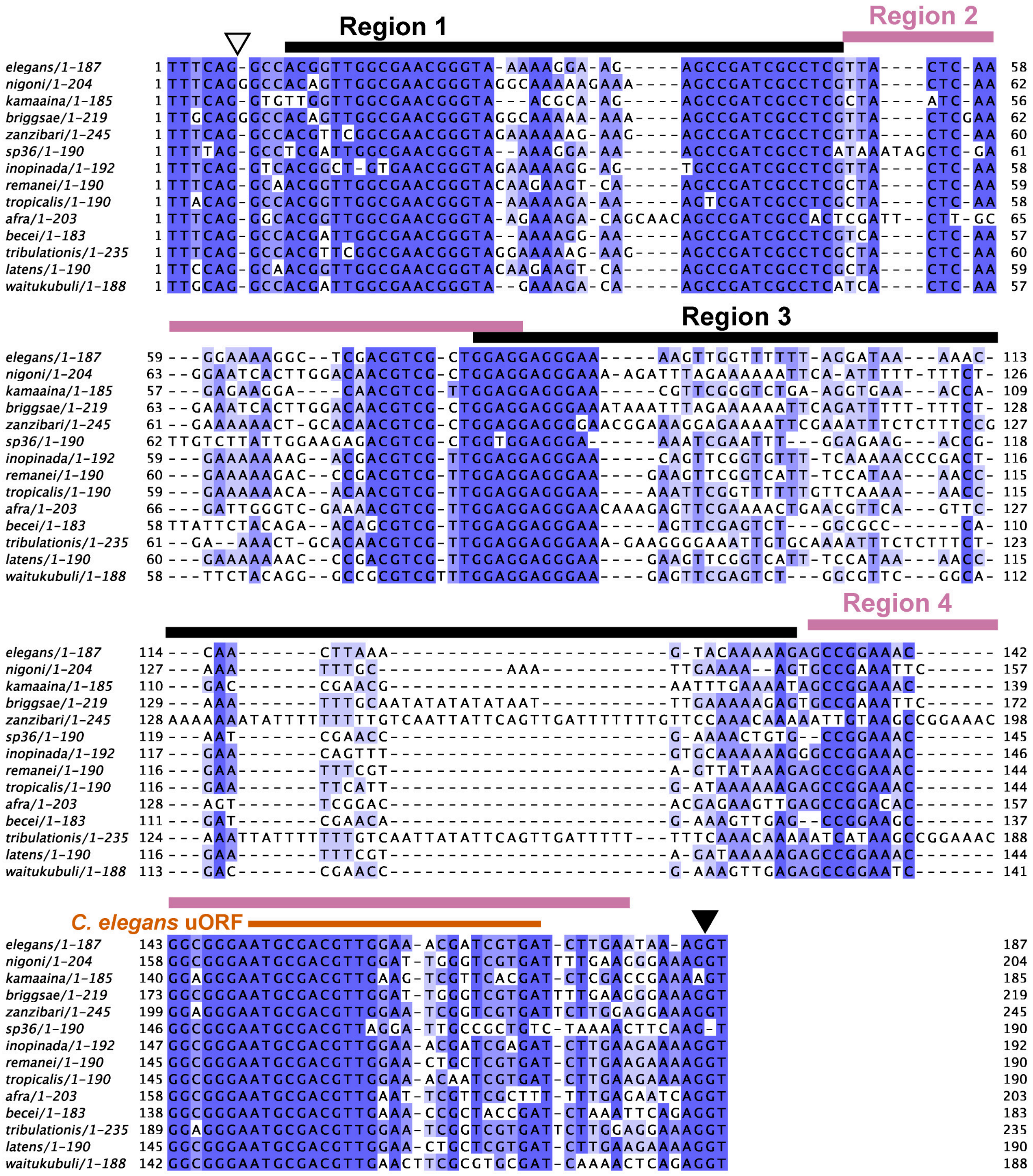
Region 1
Region 2
Region 3
Region 4
C. elegans uORF
elegans/1-187
nigoni/1-204
kamaaina/1-185
briggsae/1-219
zanzibari/1-245
sp36/1-190
inopinada/1-192
remanei/1-190
tropicalis/1-190
afra/1-203
becei/1-183
tribulationis/1-235
latens/1-190
waitukubuli/1-188

Fig. S11

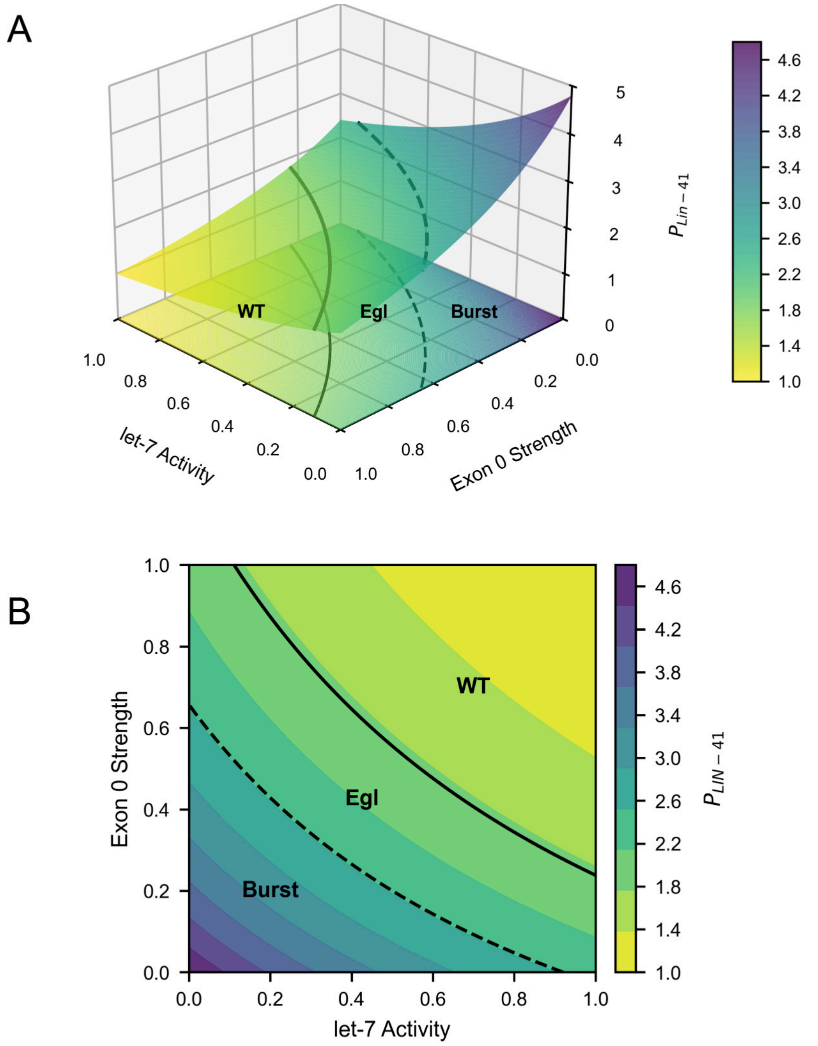
A
WT
Egl
Burst
let-7 Activity
Exon 0 Strength
$P_{Lin-41}$
B
WT
Egl
Burst
Exon 0 Strength
let-7 Activity
$P_{LIN-41}$
